# Suppressed prefrontal neuronal firing variability and impaired social representation in IRSp53-mutant mice

**DOI:** 10.1101/2021.11.17.468945

**Authors:** Woohyun Kim, Young Woo Noh, Seungjoon Lee, Woochul Choi, Se-Bum Paik, Min Whan Jung, Eunee Lee, Eunjoon Kim

## Abstract

Social deficit is a major feature of neuropsychiatric disorders, including autism spectrum disorders, schizophrenia, and attention-deficit/hyperactivity disorder, but its neural mechanisms remain unclear. Here, we examined neuronal discharge characteristics in the medial prefrontal cortex (mPFC) of IRSp53-mutant mice, which show social deficits, during social approach. IRSp53-mutant excitatory mPFC neurons displayed an increase in baseline neuronal firing and decreases in variability and dynamic range of firing rates and burst firing during social and non-social target approaches compared to wild-type controls. As a consequence, their firing activity was less differential between social and non-social targets. In addition, there was a decrease in the proportion of excitatory mPFC neurons encoding social information but not that of those encoding non-social information. These results suggest that insufficient neuronal activity dynamics may underlie impaired cortical encoding of social information and social behaviors in IRSp53-mutant mice.

## Introduction

Social dysfunction is a key feature of various neuropsychiatric disorders, including autism spectrum disorders (ASD), schizophrenia, and attention-deficit/hyperactivity disorders (ADHD). Among the various brain regions involved in social regulation, the medial prefrontal cortex (mPFC) plays critical roles in integrative and higher cognitive brain functions (Yan and Rein, 2021; Yizhar and Levy, 2021). Previous studies identified a number of mechanisms associated with dysfunctions under social context. Examples include imbalance of neuronal excitation/inhibition (Selimbeyoglu et al., 2017; Yizhar et al., 2011) (reviewed in (Lee et al., 2017; Nelson and Valakh, 2015; Sohal and Rubenstein, 2019)), impaired cortical social representation (Lee et al., 2021a; Lee et al., 2021b; Lee et al., 2016; Levy et al., 2019; Miura et al., 2020), and disruption of local oscillations (Cao et al., 2018b; Yizhar et al., 2011). Given that social behaviors represent outcomes of complex interactions among multiple underlying neural processes, further mechanistic explorations are needed to investigate such functions in the context of additional genes and various psychiatric disorders.

Insulin receptor substrate protein 53 kDa (IRSp53) encoded by the BAIAP2 gene is a postsynaptic scaffolding and adaptor protein at excitatory synapses that interacts with other key components of the postsynaptic density such as PSD-95 (Choi et al., 2005; Soltau et al., 2004). IRSp53 has also been implicated in ASD (Toma et al., 2011), schizophrenia (Fromer et al., 2014) and ADHD (Ribases et al., 2009). Functionally, IRSp53 regulates actin filament dynamics at excitatory synapses and dendritic spines (Kang et al., 2016; Scita et al., 2008).

IRSp53 deficiency in mice leads to excitatory synaptic deficits and various behavioral deficits, including hyperactivity, cognitive impairments, and social deficits (Bobsin and Kreienkamp, 2016; Chung et al., 2015; Kim et al., 2009; Kim et al., 2020; Sawallisch et al., 2009). IRSp53 knockout (KO) mice have fewer dendritic spines and enhanced NMDA receptor (NMDAR) function; they show impaired social behavior that is rescued by pharmacological NMDAR suppression (Chung et al., 2015; Kim et al., 2009). Importantly, mPFC neurons in IRSp53-KO mice show reduced neuronal firing under urethane-anesthesia, which is acutely normalized by pharmacological NMDAR suppression (Chung et al., 2015). However, it remained unknown whether and how the social behavioral deficits are associated with altered mPFC neural activity in waking-state animals engaged in social interaction.

To study the neural abnormalities of the mPFC associated with social dysfunction in IRSp53-KO mice, we herein performed single-unit recording in freely moving mice engaged in social interaction in a linear social-interaction chamber (Lee et al., 2016). We found that excitatory neurons in the mPFC of IRSp53-KO mice display narrower dynamic ranges of firing rate and lower discrimination between social and object targets compared to those of wild-type (WT) controls. Our results uncover a novel social coding deficit associated with IRSp53-KO.

## Results

### Social impairments in IRSp53-KO mice in the linear-chamber social-interaction test

To compare neuronal activities in the mPFC of WT and IRSp53-KO mice during social interaction, we performed single-unit recordings in mice engaged in social interaction in a linear-chamber social-interaction apparatus (**Figure 1A**). The chamber, a long corridor connected with two side chambers with targets, was designed to measure mPFC activity during social interaction (Lee et al., 2016). A subject mouse was first placed in a separate rest box (7.5 x 15 cm) for 5 minutes for recording of resting neural activity. The mouse was then placed into the linear social-interaction chamber and allowed to explore the chamber with both side chambers being empty (empty-empty/E-E session) for 10 minutes. This was followed by a session in which one of the side chambers contained a novel social target (S; a conspecific male mouse) and the other contained a novel inanimate object (O) (first S-O session), and another session where S and O were switched (second S-O session), which was included to control for side (or location)-specific as opposed to target-specific neural activity. The positions of mice in the linear chamber during experiments were determined using the DeepLabCut program (Mathis et al., 2018), which automatically marked the mouse’s nose, ears, and tail base (**Figure 1B**). Sniffing time was defined as the time when the mouse’s nose was within a distance of 3 cm from the front face of the target chamber. In-zone time was defined as the time when the body center (midpoint between the nose and tail base) fell in the area within 9 cm from the front face of the target chamber.

**Figure 1.**
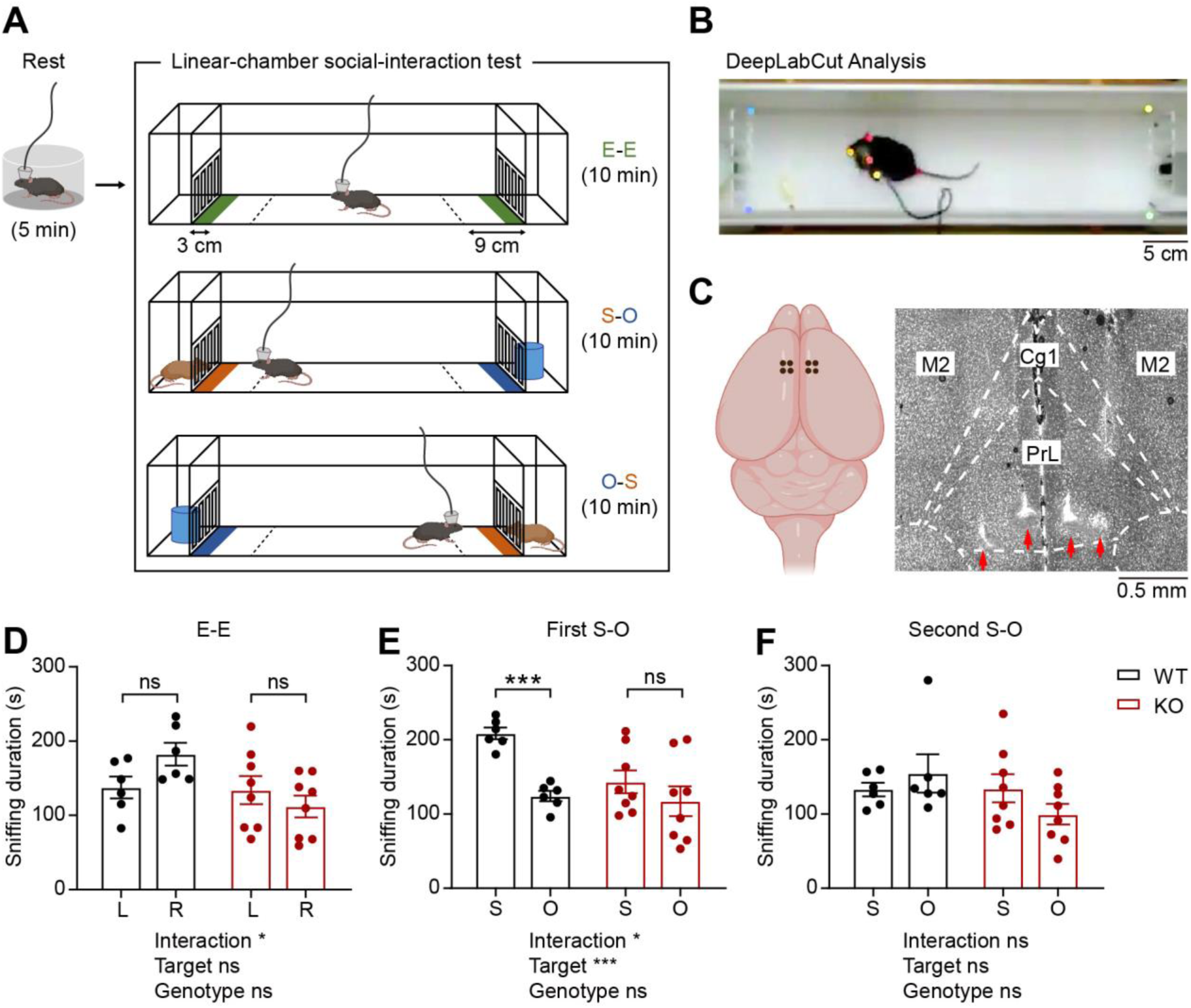
Social impairments in IRSp53-KO mice in the linear-chamber social-interaction test. **(A)** Schematic diagram of the linear-chamber social-interaction test used to measure social approach towards a novel conspecific mouse (S, social) versus a novel non-social target (O, object). A tetrode-implanted mouse was first placed in the rest box for 5 minutes and moved to the linear social-interaction chamber to perform the following three sessions: empty-empty (E-E) session, social-object (first S-O) session, and object-social (second S-O) session. The in-zone areas, falling within 9 cm from the front faces of the chambers, are indicated by the dashed lines. The sniffing zones, falling within 3 cm from the front faces of the chambers, are indicated by green, orange, and blue colors. **(B)** An example video frame of mouse body parts automatically tracked by the DeepLabCut program. **(C)** Schematic (left) and a representative coronal brain section (right) showing the locations of the implanted tetrodes. PrL, prelimbic cortex; IL, infralimbic cortex; Cg1, cingulate cortex, area 1; M2, secondary motor cortex. **(D–F)** Mean sniffing durations (±standard error of mean/SEM) for left (L) vs. right (R) empty targets during the E-E session (**D**) and the social (S) vs. object (O) targets during the first S-O (**E**) and second S-O (**F**) sessions. (n = 6 mice [WT], 8 mice [IRSp53-KO], *p < 0.05, ***p < 0.001, ns, not significant, two-way repeated-measures (RM)-ANOVA with Sidak’s multiple comparisons test). See **Supplementary file 2** for statistics. Numerical data used to generate the figure are available in the **Figure 1—source data 1**. Figure 1—source data 1 **Source files for mouse behavior data in Figure 1** The excel file contains the numberical data used to generate Figure 1D–F.

The single-unit activity was recorded with tetrodes from the prelimbic (PrL), infralimbic (IL), and cingulate cortex (Cg1) regions. Eight tetrodes, four tetrodes in each hemisphere, were implanted into the mPFC and lowered after each round of recording experiment to record neurons at different depths. After the last recording, the locations of all tetrodes were assessed via histology, and data from those falling within the area of interest were used for analysis (**Figure 1C, Figure 1—figure supplement 1A**).

In the E-E session, WT and IRSp53-KO mice did not show preference to the left- or right-side chamber, as assessed by sniffing and in-zone durations (**Figure 1D, Figure 1—figure supplement 2A**). In the first S-O session, IRSp53-KO mice spent a comparable amount of time exploring the social and object targets, whereas WT mice displayed a strong preference for the social target (**Figure 1E, Figure 1— figure supplement 2B**). In the second S-O session, WT mice no longer displayed social preference, likely because of social habituation (**Figure 1F, Figure 1—figure supplement 2C**).

While IRSp53-KO mice showed decreased sniffing visits to the social conspecific mouse, their mean duration of each visit was comparable to that of the WT mice (**Figure 1—figure supplement 2D, E**). Moreover, there was no genotype difference in the total distance travelled (**Figure 1—figure supplement 2F, G**). WT and IRSp53-KO mice displayed a decline in the locomotor activity across successive sessions (E-E, first S-O, and second S-O) in each recording experiment (**Figure 1— figure supplement 2F**), but their overall locomotion remained comparable across the ten experiments (**Figure 1—figure supplement 2G**). These results collectively indicate that IRSp53-KO mice display social impairments in the linear social-interaction chamber, similar to the social impairments previously observed in three-chamber and direct/dyadic social-interaction tests (Chung et al., 2015).

### Increased resting firing rate in IRSp53-KO pExc mPFC neurons

We next compared neuronal firing patterns in the mPFC of WT and IRSp53-KO mice during the abovementioned linear-chamber social-interaction test. To this end, we first analyzed rest-period firing rates in awake and freely moving WT and IRSp53-KO mice. We segregated the neurons into putative excitatory (pExc) and putative inhibitory (pInh) neurons based on their half-valley width (pExc > 200 ms; pInh < 200 ms) and peak-to-valley ratio (pExc > 1.4; pInh < 1.4) (**Figure 2A, B**). The firing rate of total neurons at rest was increased in the mPFC of IRSp53-KO mice, compared with WT mice (**Figure 2C**). However, only the IRSp53-KO pExc neurons, but not IRSp53-KO pInh neurons, showed a significant increase in firing rate (**Figure 2D, E**), suggesting that pExc neurons mainly contribute to the increase in the total firing rate. These results differ from those previously obtained from anesthetized IRSp53-KO mice (Chung et al., 2015), which exhibited decreases in total and pExc firing. This highlights the importance of measuring cortical neuronal activity in behaving mice engaged in social interaction.

**Figure 2.**
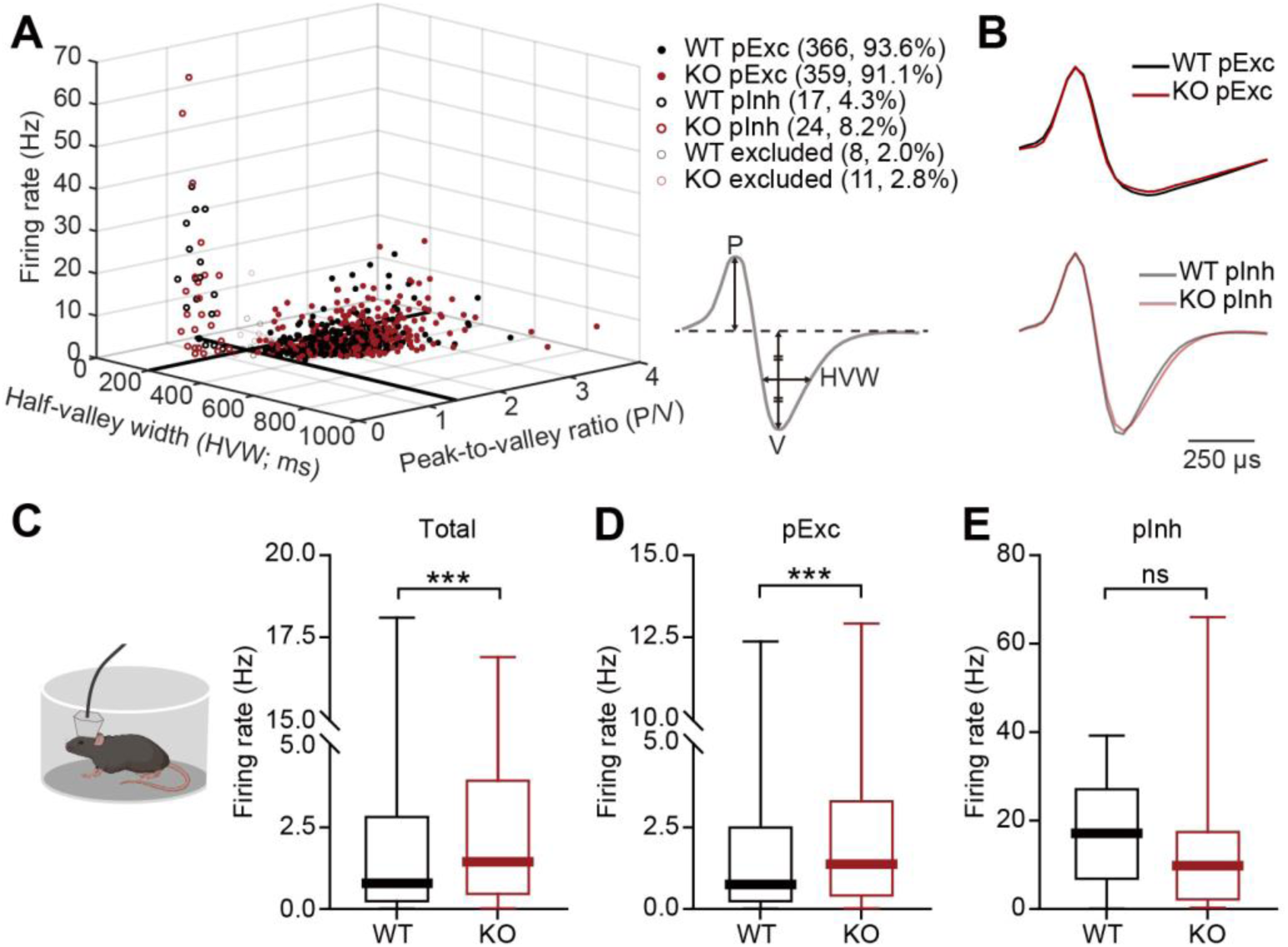
Increased resting firing rate in IRSp53-KO pExc mPFC neurons. **(A)** Classification of recorded neurons into putative excitatory (pExc) and putative inhibitory (pInh) neurons based on the half-valley width (200 ms) and peak-to-valley ratio (1.4). P, peak; V, valley; HVW, half-valley width. **(B)** Average waveforms of WT and IRSp53-KO pExc (top) and pInh (bottom) neurons. The waveform of each neuron was normalized by its peak value. **(C–E)** Firing rates of WT and IRSp53-KO total (**C**), pExc (**D**), and pInh (**E**) neurons in the mPFC during the 5-min rest period. (n = 391 [WT-total], 394 [KO-total], 366 [WT-pExc], 359 [KO-pExc], 17 [WT-pInh], 24 [KO-pInh], ***p < 0.001, ns, not significant, Mann-Whitney test). See **Supplementary file 2** for statistics. Numerical data used to generate the figure are available in the **Figure 2—source data 1**. Figure 2—source data 1 **Source files for resting firing rate data in Figure 2** The excel file contains the numberical data used to generate Figure 2A–E.

It should be noted that the majority of recorded neurons were pExc neurons (WT: 366 neurons, 93.6%, IRSp53-KO: 359 neurons, 91.1%), and that relatively few recordings were obtained from pInh neurons (WT: 17 neurons, 4.3%, IRSp53-KO: 24 neurons, 8.2%). Because IRSp53 is expressed primarily in the excitatory (not inhibitory) pyramidal neurons of the cortex (Burette et al., 2014), we hypothesized that the main effects of IRSp53 loss are seen in the pExc neurons. Therefore, only pExc neurons were used for further analysis. Of all pExc neurons recorded, only those with a mean firing rate of ≥ 0.5 Hz were included for further analysis in order to avoid low sampling errors arising from the inclusion of neurons with low firing rates (Supplementary file 1).

### Reduced firing-rate range and variability in IRSp53-KO pExc mPFC neurons

The pExc mPFC neurons of IRSp53-KO mice showed significantly higher mean firing rates than those of WT mice during the initial 5-min rest period, but not during the 30-min linear chamber test period (**Figure 3B**). Thus, pExc neurons of the mPFC did not differ significantly between IRSp53-KO and WT mice in terms of overall mean firing rates during the linear chamber test. However, we noticed in our preliminary analysis that the temporal profiles of instantaneous firing rates (3-sec time-bin advanced in 1-sec steps) differ substantially between WT and IRSp53-KO pExc neurons during the 30-min linear chamber test, as shown by the representative examples in **Figure 3A**.

**Figure 3.**
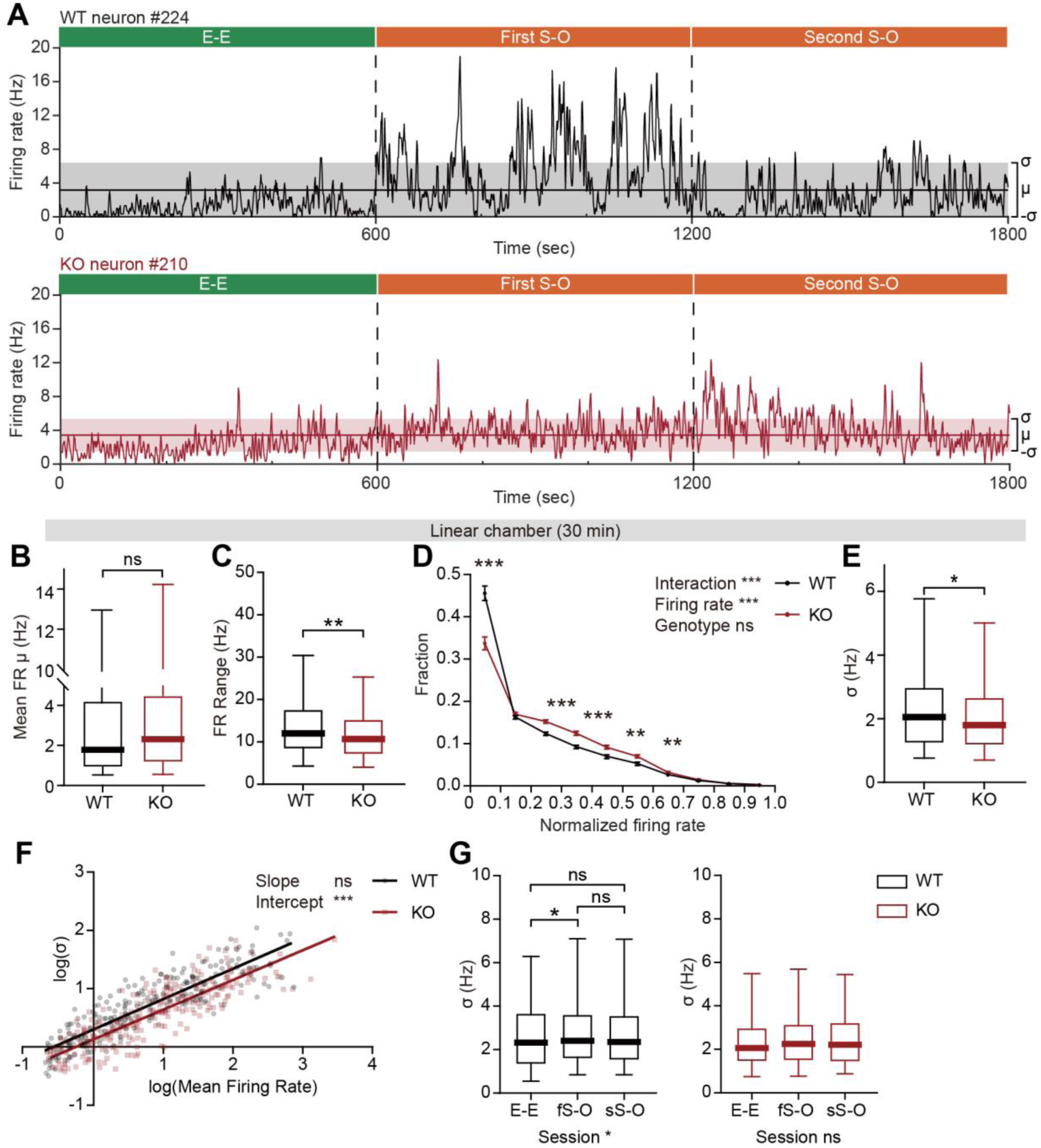
Decreased firing-rate range and variability in IRSp53-KO pExc mPFC neurons during linear chamber exploration. **(A)** Instantaneous firing-rate traces of representative WT (top) and IRSp53-KO (bottom) pExc neurons (3-sec window advanced in 1-sec steps) during a sample linear chamber experiment (30 min). Solid horizontal lines indicate mean firing rates (μ). Shaded regions indicate one standard deviation (σ, sigma). **(B)** Mean firing rates of WT and IRSp53-KO pExc neurons during the 30-min linear chamber test. (n = 233 [WT-pExc] and 258 [KO-pExc], ns, not significant, Mann-Whitney test). **(C)** Firing-rate ranges (maximum – minimum instantaneous firing rate) of WT and IRSp53-KO pExc neurons during the linear chamber test (n = 233 [WT-pExc] and 258 [KO-pExc], **p < 0.01, Mann-Whitney test). **(D)** Mean (±SEM) histograms of normalized instantaneous firing rate during the linear chamber test. For each neuron, instantaneous firing rates were normalized by its maximum instantaneous firing rates. (n = 233 [WT-pExc] and 258 [KO-pExc], **p < 0.01, ***p < 0.001, ns, not significant, two-way RM-ANOVA with Bonferroni’s multiple comparisons test). **(E)** Sigma values of the instantaneous firing rates of WT and IRSp53-KO pExc neurons during the linear chamber test. (n = 233 [WT-pExc] and 258 [KO-pExc], *p < 0.05, Mann-Whitney test). **(F)** Log-scale scatter plot of sigma values against mean firing rates of WT and IRSp53-KO pExc neurons during the linear chamber test. Solid lines indicate simple linear regression of WT (black) and KO (red) values. (n = 233 [WT-pExc] and 258 [KO-pExc], ***p < 0.001, ns, not significant, slope comparison test (see Methods)). **(G)** Sigma values for the instantaneous firing rates of WT (left) and IRSp53-KO (right) pExc neurons during the E-E, first S-O, and second S-O sessions of the linear chamber test. (n = 233 [WT-pExc] and 258 [KO-pExc], *p < 0.05, ns, not significant, Friedman test followed by Dunn’s multiple comparisons test). See **Supplementary file 2** for statistics. Numerical data used to generate the figure are available in the **Figure 3—source data 1**. Figure 3—source data 1 **Source files for instantaneous firing rate data in Figure 3** The excel file contains the numberical data used to generate Figure 3A–G.

Further examinations of instantaneous firing rate revealed that the maximum instantaneous firing rate during the linear chamber test was significantly lower in IRSp53-KO pExc neurons than WT pExc neurons. In contrast, there was a trend for higher minimum instantaneous firing rates in IRSp53-KO pExc neurons than WT pExc neurons (*p* = 0.0544; **Figure 3—figure supplement 1A,B**). Consequently, the dynamic range of firing rate (the difference between the maximum and minimum instantaneous firing rates) during the linear chamber test was significantly narrower for IRSp53-KO pExc neurons than WT pExc neurons (**Figure 3C**).

Another difference we noticed was that while WT neurons often remain silent and show abrupt increases in firing rate at specific time points, IRSp53-KO neurons tended to be active more chronically with their instantaneous firing rates fluctuating around the mean (**Figure 3A**). To test whether this is indeed the case, we examined the distribution of instantaneous firing rates of WT and IRSp53-KO neurons (normalized to the maximum firing rate). As expected, IRSp53-KO neurons had a significantly lower proportion of time-bins in the lowest firing rate (0–0.1) and instead higher proportions in mid-range firing rates (0.2–0.6) compared to the WT neurons (**Figure 3D**).

Given the reduced firing-rate range, we speculated that the firing rate variability of IRSp53-KO pExc neurons may also be decreased. We defined the firing rate variability of each neuron by the sigma value (1 standard deviation around the mean) of its instantaneous firing rates. We found that the instantaneous firing rates of IRSp53-KO neurons were indeed less variable, as indicated by a decrease in the sigma value (**Figure 3E**). This decrease in sigma value was observed consistently across the analyses using variable sizes of time window for calculating instantaneous firing rate, ranging from 0.5 to 5 s (**Figure 3—figure supplement 1C**). This phenomenon was specific to the recordings from the linear chamber sessions but not the rest period (**Figure 3—figure supplement 1D**). In order to test if this decrease in the variability in the IRSp53-KO neurons is dependent upon the mean firing rate, the relationship between mean firing rates and sigma values was compared between genotypes (**Figure 3F**). As indicated by the significant decrease in the intercept—and comparable slope— of the linear regression, IRSp53-KO neurons were generally less variable in instantaneous firing activity regardless of their mean discharge rate.

We next examined whether firing rate variability varies across the three sessions (E-E, first S-O, and second S-O) of the linear chamber test. We found that WT neurons showed increased variability in instantaneous firing rate during the first S-O session compared to the E-E session. In contrast, IRSp53-KO neurons showed similar levels of variability across the three sessions (**Figure 3G**). The increase in the variability of IRSp53-KO neuronal activity during the first S-O session could not be accounted for by the difference in mean firing rate (**Figure 3—figure supplement 1E**). These results collectively indicate that excitatory mPFC neurons in IRSp53-KO mice have reduced firing-rate range and variability.

### Impaired bursting in IRSp53-KO pExc mPFC neurons

Based on the reduction in the firing-rate range, we reasoned that there may be a shift in the distribution of interspike intervals (ISIs) in IRSp53-KO pExc neurons. Contrary to the comparable levels of average ISI histograms between WT and IRSp53-KO pExc neurons at rest, those during the linear chamber test differed in IRSp53-KO pExc neurons. In particular, there was a pronounced reduction in the proportion of ISIs ≤ 10 ms (**Figure 4A, B**). This result suggests that the ability to exhibit an abrupt increase in firing rate may be impaired in IRSp53-KO pExc neurons.

**Figure 4.**
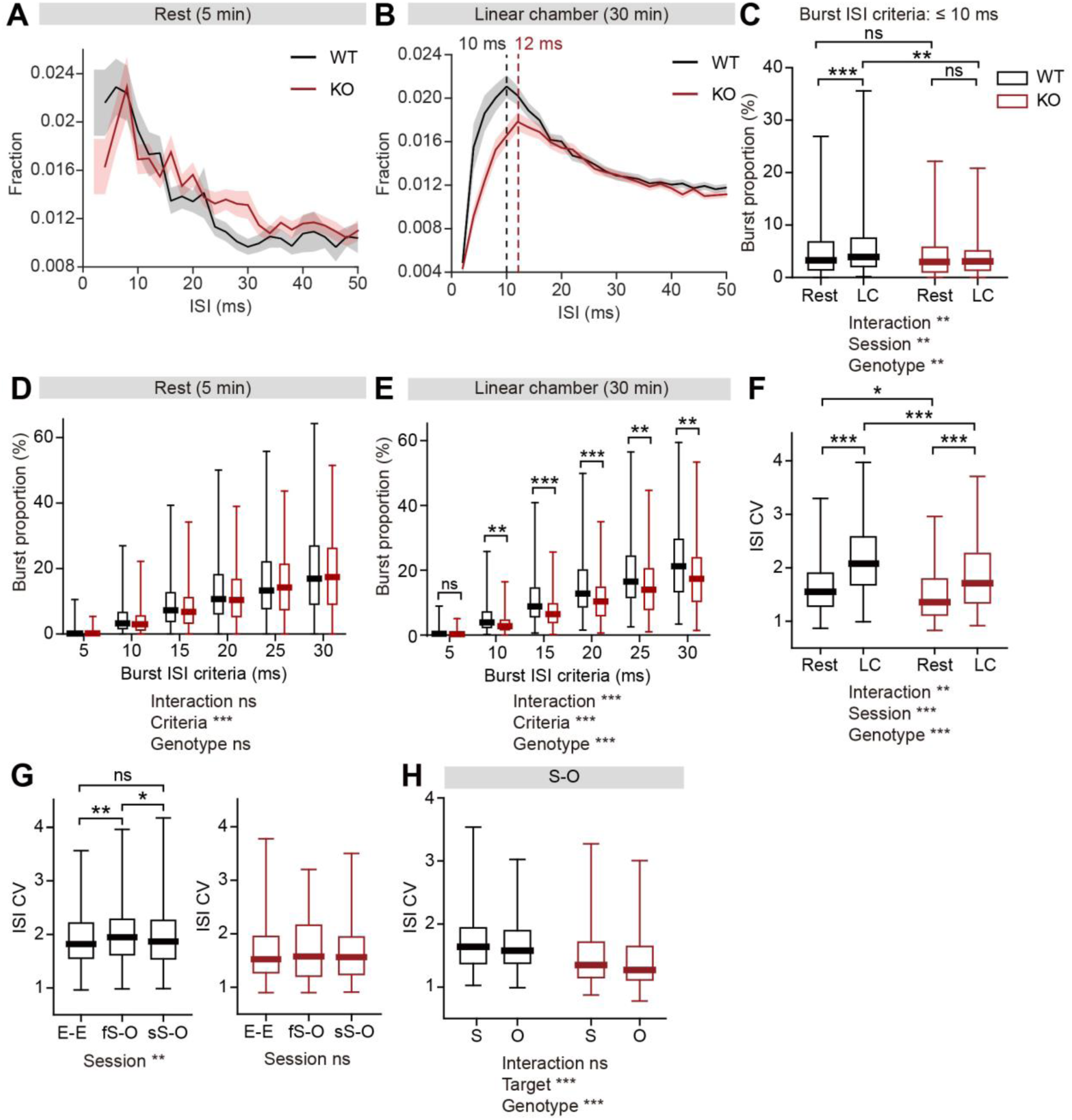
Lower burst firing and spike variability in IRSp53-KO pExc mPFC neurons during linear chamber exploration. **(A and B)** Mean (±SEM) histograms of interspike intervals (ISI) at the 5-min rest (**A**) and the 30-min linear chamber (**B**) periods. Note that the peaks of mean ISI distributions for WT (black) and IRSp53-KO (red) neurons fall at 10 ms and 12 ms, respectively (indicated by dashed lines). (n = 233 [WT-pExc] and 258 [KO-pExc]). **(C)** Burst proportions (proportion of burst spikes out of total spikes) of WT and IRSp53-KO pExc neurons during the rest and linear chamber (LC) periods for burst ISI threshold of 10 ms. (n = 233 [WT-pExc] and 258 [KO-pExc], **p < 0.01, ***p < 0.001, ns, not significant, two-way RM-ANOVA with Sidak’s multiple comparisons test). **(D and E)** Burst proportions of WT and IRSp53-KO pExc neurons during the rest (**D**) and linear chamber (**E**) periods, for different burst ISI thresholds. (n = 233 [WT-pExc] and 258 [KO-pExc], *p < 0.05, **p < 0.01, ***p < 0.001, ns, not significant, two-way RM-ANOVA with Sidak’s multiple comparisons test). **(F)** Coefficient of variations (CVs) of ISIs for WT and IRSp53-KO pExc neurons during the rest and linear chamber periods. (n = 233 [WT-pExc] and 258 [KO-pExc], *p < 0.05, **p < 0.01, ***p < 0.001, two-way RM-ANOVA with Sidak’s multiple comparisons test). **(G)** ISI CVs for WT (left) and IRSp53-KO (right) pExc neurons during the E-E, first S-O, and second S-O sessions of the linear chamber test. (n = 233 [WT-pExc] and 258 [KO-pExc], *p < 0.05, **p < 0.01, ns, not significant, Friedman test with Dunn’s multiple comparisons test). **(H)** ISI CVs for WT and IRSp53-KO pExc neurons during social (S) versus object (O) sniffing in the first and second S-O sessions combined. (n = 233 [WT-pExc] and 258 [KO-pExc], ***p < 0.001, ns, not significant, two-way RM-ANOVA with Sidak’s multiple comparisons test). See **Supplementary file 2** for statistics. Numerical data used to generate the figure are available in the **Figure 4—source data 1**. Figure 4—source data 1 Source files for ISI and burst data in Figure 4 The excel file contains the numberical data used to generate Figure 4A–H.

Because there was a shift in the ISI distribution of IRSp53-KO pExc neurons, we reasoned that burst firing might be reduced in IRSp53-KO mice. We defined burst spikes as those with short ISIs (≤ 10 ms) during linear chamber exploration (**Figure 4B**). Comparing burst firing across the rest and linear chamber periods, we found that burst firing (ISI ≤ 10 ms) increased significantly when WT mice switched from the resting state to linear chamber-exploring state. Such change, however, was not observed in IRSp53-KO mice (**Figure 4C**). Burst firing did not differ significantly between WT and IRSp53-KO pExc neurons during the rest period, but was significantly lower in IRSp53-KO pExc neurons than WT pExc neurons during the linear chamber sessions (**Figure 4C**).The same conclusion was obtained when we increased the cut-off value for burst spikes up to 30 ms (**Figure 4D, E**).

The majority of burst events for pExc neurons (∼70–90%) were spike doublets (two-spike events) in both WT and IRSp53-KO mice. However, the proportions of burst events with three spikes (spike triplets) or ≥4 spikes were significantly lower in IRSp53-KO neurons than WT neurons (**Figure 4—figure supplement 1A**). In addition, the composition of burst events (according to spike count) varied between social and object targets in WT, but not IRSp53-KO, pExc neurons (**Figure 4—figure supplement 1B, C**). WT burst events during social target sniffing consisted of significantly higher proportions of triplet and ≥4 spikes burst events, compared to those during object target sniffing, in the analysis using moderate cut-off values for burst spikes (15–30 ms) (**Figure 4—figure supplement 1B, C**).

The coefficient of variation (CV) of ISIs is a well-known measure of single neuronal spike variability (Sendhilnathan et al., 2020). Confirming the results obtained from the analysis using sigma values, we found that the CV of ISIs was significantly lower for IRSp53-KO pExc neurons compared to WT pExc neurons. Moreover, CV of ISIs increased during linear chamber exploration relative to the rest period in both genotypes (**Figure 4F**). Consistent with the results obtained from the analysis of sigma value, WT neurons, but not IRSp53-KO neurons, showed a significant increase in spike variability during the first S-O session compared to E-E session (**Figure 4G**). Interestingly, ISI CVs were generally higher in both WT and IRSp53-KO pExc neurons during social target sniffing compared to object target sniffing; however, the overall ISI CVs were lower in IRSp53-KO pExc neurons (**Figure 4H**).

Taken together, these results suggest that IRSp53-KO pExc neurons show diminished burst firing and a weakened ability to discriminate social and object targets by distinct burst event compositions.

### Weak responses to social and object targets in IRSp53-KO pExc mPFC neurons

In the linear chamber, mice can actively explore the targets or spend time in non-target areas. We reasoned that, if the firing-rate range of IRSp53-KO mice is decreased, the magnitude of neuronal responses to the social or object stimulus may also be decreased. To test this, we divided the linear chamber into five equal-area sections (each 9-cm in length), and the firing rates in the in-zone areas were compared to that of the center zone (**Figure 5A**). We defined the maximum Δ firing rate of each session as the maximum absolute difference in mean firing rates between the center zone (FR_c_) and the two in-zones (FR_I1_, FR_I2_; left and right for E-E session, social and object for S-O sessions).

**Figure 5.**
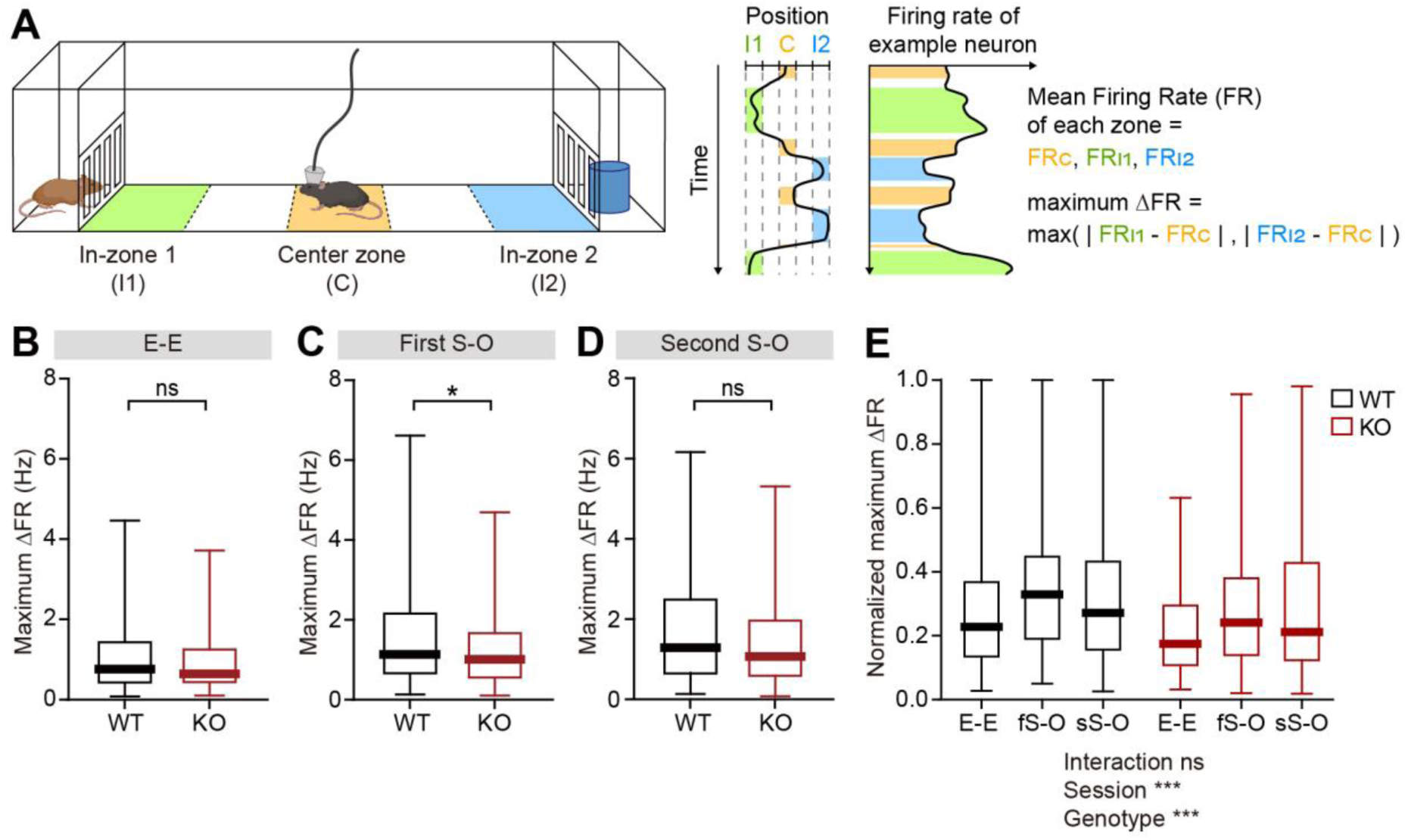
Limited firing-rate changes in response to social and object targets in IRSp53-KO pExc mPFC neurons. **(A)** Definition of in-zone and center zone. The first and fifth of the equally divided five 9-cm-long areas were defined as in-zones (I1 and I2, respectively) while the third area was defined as the center zone (C). For each neuron, the maximum Δ firing rate is defined as the higher value among the firing rate differences between the center zone and two in-zones (left and right for E-E session, social and object for S-O sessions). **(B–D)** Maximum Δ firing rates of WT and IRSp53-KO pExc neurons during the E-E (**B**), first S-O (**C**), and second S-O (**D**) sessions. (n = 233 [WT-pExc] and 258 [KO-pExc], *p < 0.05, ns, not significant, Mann-Whitney test). **(E)** Normalized maximum Δ firing rates of WT and IRSp53-KO pExc neurons during the E-E, first S-O (fS-O), and second S-O sessions (sS-O). (n = 233 [WT-pExc] and 258 [KO-pExc], ***p < 0.001, ns, not significant, two-way RM-ANOVA). See **Supplementary file 2** for statistics. Numerical data used to generate the figure are available in the **Figure 5—source data 1**. Figure 5—source data 1 Source files for firing-rate change data in Figure 5 The excel file contains the numberical data used to generate Figure 5B–E.

Compared to WT pExc neurons, IRSp53-KO pExc neurons displayed a decreased maximum Δ firing rate only in the first S-O session of the linear-chamber test, but not in the E-E or second S-O session (**Figure 5B–D**). IRSp53-KO pExc neurons showed a general decrease in the normalized maximum Δ firing rate (see Methods) across all three sessions (**Figure 5E**). It is notable that the response magnitudes of both WT and IRSp53-KO pExc neurons were the highest during the first S-O session, in response to novel social and object targets. This fits well with our behavioral data, in which only WT mice, but not IRSp53-KO mice, show social preference in the first, but not second, S-O session (**Figure 1D–F, Figure 1—figure supplement 2A–C**).

### Limited social versus object firing-rate discriminability in IRSp53-KO pExc mPFC neurons

Since the firing-rate range of IRSp53-KO neurons was found to be limited, especially in the first S-O session, we hypothesized that the firing-rate discriminability between social and object targets may also be limited. The slopes of linear regression and the degrees of dispersion (indicated by 95% confidence interval) for the left versus right (L vs. R) in-zone firing rates in the E-E session were comparable between genotypes (**Figure 6A**). In contrast, the slopes of the linear regression lines relating social and object (S vs. O) firing rates were biased towards the social firing rate in both genotypes in the first and second S-O sessions, indicating preferential responses to social to object targets. However, this bias was smaller in IRSp53-KO pExc neurons compared to WT pExc neurons (**Figure 6B, Figure 6—figure supplement 1A**). Additionally, the confidence interval was narrower for IRSp53-KO pExc neurons, especially in the first S-O session, compared to WT pExc neurons for the S versus O firing rates, suggesting that the former exhibited limited discriminability (**Figure 6B**). Consistently, the absolute difference in the firing rate for S versus O (an indication of discriminability) in the first S-O session was significantly lower in IRSp53-KO pExc neurons than WT pExc neurons (**Figure 6C**). Nevertheless, IRSp53-KO pExc neurons could still discriminate the social targets from object targets significantly better compared to the left versus right side discrimination in the E-E session (**Figure 6C**). This result suggests that IRSp53-KO mice have the ability to recognize social and object stimuli, albeit in reduced degree than WT mice.

**Figure 6.**
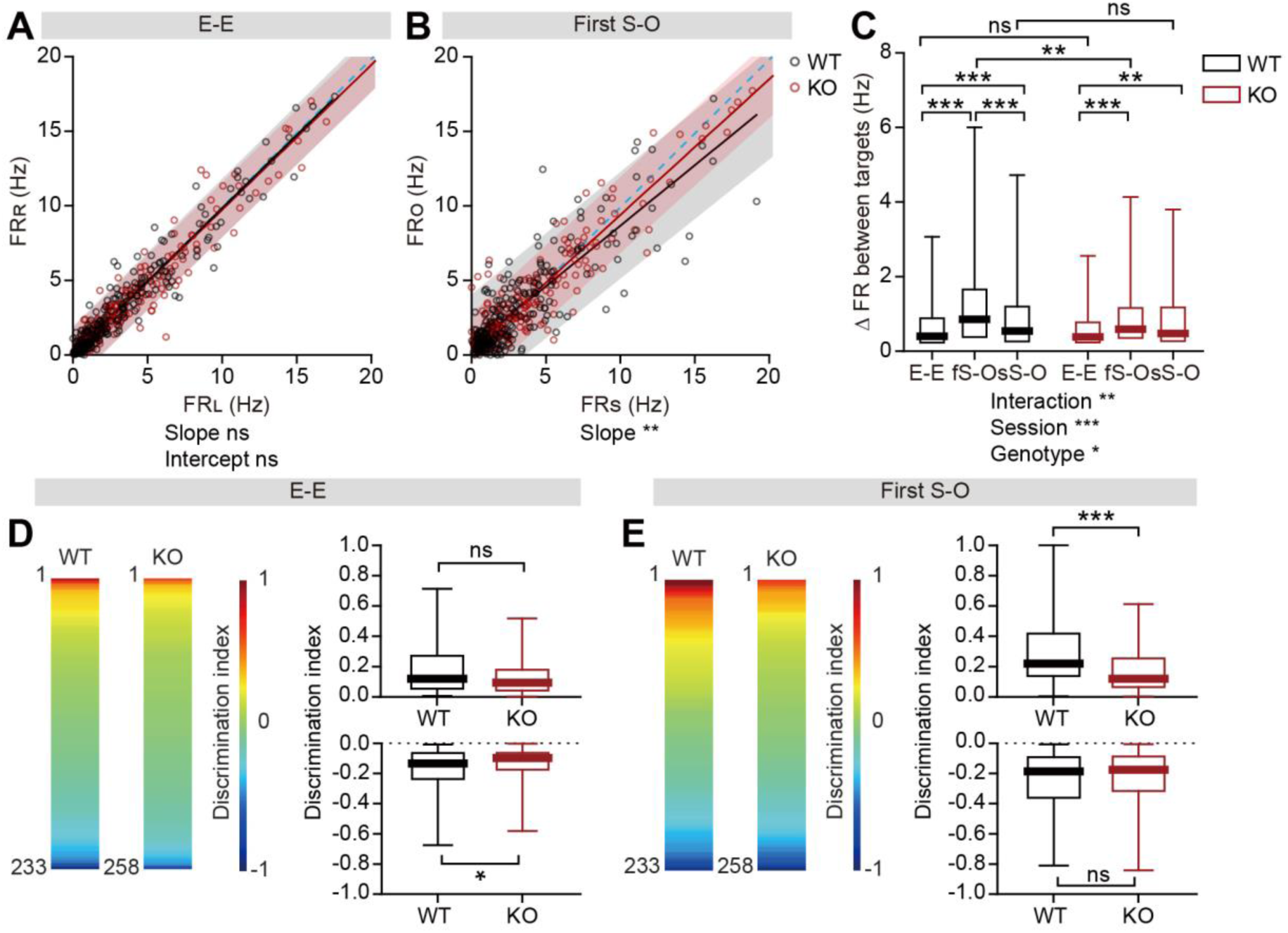
Lower firing-rate discriminability between social and object targets in IRSp53-KO pExc mPFC neurons. **(A and B)** Scatterplots of left in-zone firing rate (FR_L_) against right in-zone firing rate (FR_R_) during the E-E session (**A**) and social in-zone firing rate (FR_S_) against object in-zone firing rate (FR_O_) during the first S-O session (**B**) for WT and IRSp53-KO pExc neurons. Solid lines indicate simple linear regressions for WT (black) and KO (red) neurons. Shaded areas indicate the 95% confidence intervals for the WT (black) and KO (red) firing rates. Blue dashed lines are 45-degree lines. (n = 233 [WT-pExc] and 258 [KO-pExc], **p < 0.01, ns, not significant, simple linear regression with slope comparison test (see Methods)). **(C)** Absolute changes in left versus right in-zone firing rates (E-E session) and social versus object in-zone firing rates (first (fS-O) and second (sS-O) S-O sessions) for WT and IRSp53-KO pExc neurons. (n = 233 [WT-pExc] and 258 [KO-pExc]), *p < 0.05, **p < 0.01, ***p < 0.001, ns, not significant, two-way RM-ANOVA with Sidak’s multiple comparisons test). **(D)** Heatmaps (left) showing the discrimination index representing side discriminability during the E-E session of individual WT and IRSp53-KO pExc neurons sorted from 1 to –1 (n = 233 [WT-pExc] and 258 [KO-pExc]). Positive (top, n = 104 [WT-pExc] and n = 123 [KO-pExc]) and negative (bottom, n = 129 [WT-pExc] and n = 135 [KO-pExc]) discrimination indexes (right) represent pExc neurons with left > right and left < right discriminability, respectively, during the E-E session. (*p < 0.05, ns, not significant, Mann-Whitney test). **(E)** Heatmaps (left) showing the discrimination index representing social vs. object target discriminability during the first S-O session of individual WT and IRSp53-KO pExc neurons sorted from 1 to –1 (n = 233 [WT-pExc] and 258 [KO-pExc]). Positive (top, n = 115 [WT-pExc] and n = 128 [KO-pExc]) and negative (bottom, n = 118 [WT-pExc] and n = 130 [KO-pExc]) discrimination indexes (right) represent pExc neurons with social > object and social < object discriminability, respectively, during the first S-O session. (***p<0.001, ns, not significant, Mann-Whitney test). See **Supplementary file 2** for statistics. Numerical data used to generate the figure are available in the **Figure 6—source data 1**. Figure 6—source data 1 Source files for firing-rate discriminability data in Figure 6 The excel file contains the numberical data used to generate Figure 6A–E.

As shown by the heatmap for the discrimination index (DI; normalized quantification of discriminability) between two targets (L vs. R or S vs. O), some neurons showed high firing rates to the left (or social) targets relative to right (or object) targets, while others showed the opposite response patterns (**Figure 6D, E, Figure 6—figure supplement 1B**). IRSp53-KO pExc neurons with preferential firing to social than object target had significantly lower discriminability relative to those of WT neurons during the first S-O session (**Figure 6E**). Such difference between two genotypes was not observed during the second S-O session.

### Decreased social-responsive neuronal proportion in the mPFC of IRSp53-KO mice

The results so far suggest that the mPFC pExc neurons of IRSp53-KO mice may be less efficient in encoding social information, potentially leading to a decline in the proportion of social target-responsive neurons in IRSp53-KO mice. We analyzed neuronal activity during the three linear chamber sessions (E-E, first S-O, and second S-O sessions) and determined empty-, social-, and object target-responsive neurons (termed empty, social, and object neurons hereafter) as those whose firing at the target sniffing zone differed from that in the center zone (|z-score| ≥ cut-off value; see Methods). Because the optimal z-score cut-off value is not known a priori, the proportions of social, object, empty neurons were calculated for a range of z-score cut-off values (0.00–2.58; p-value 0.01–1.00), and the resulting values were subject to curve fitting. We found that the fitted curves differed significantly between WT and IRSp53-KO mice for social neurons; a significantly lower proportion was classified as social neurons among IRSp53-KO pExc neurons compared to WT pExc neurons (**Figure 7A**), whereas the proportions of object neurons and empty neurons were comparable between genotypes (**Figure 7B, Figure 7—figure supplement 1A**).

**Figure 7.**
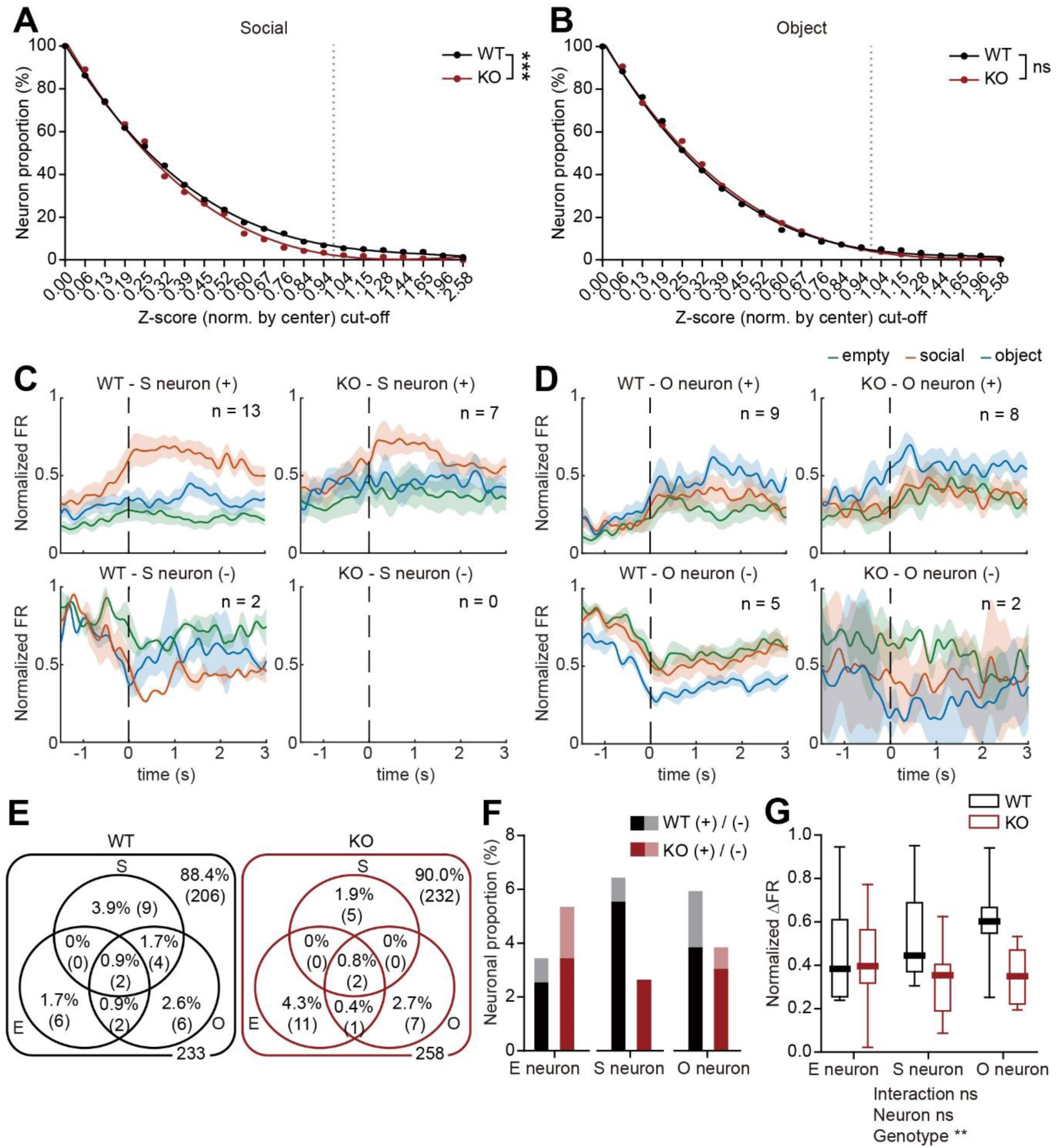
Decreased proportion of social pExc neurons in the mPFC of IRSp53 KO mice. **(A and B)** Social (**A**) and object (**B**) neuronal proportions out of 233 WT pExc neurons and 258 IRSp53-KO pExc neurons, obtained using a z-score cut-off range of 0 to 2.58. For each neuron, the mean z-scores of firing rates obtained during social and object sniffing were normalized by the firing rates obtained at the center zone. See Methods for details on z-score calculation. Solid lines indicate nonlinear fitted lines for the WT (black) and IRSp53-KO (red) groups. Dotted lines indicate z-score cut-off value of 1.0. Note that the social neuron proportion, but not the object neuron proportion, is significantly different between the genotypes. (***p < 0.001, ns, not significant, comparison of non-linear fits (see Methods)). **(C and D)** Average peristimulus time histograms (PSTHs) of firing rate responses to empty (green), social (orange), and object (blue) targets (aligned to the onset of sniffing) for all social (S; **C**) and object (O; **D**) neurons. The pExc neurons were filtered by a z-score cut-off value of 1.0. Social and object neurons are divided by genotype (WT left, IRSp53-KO right) and response direction (positive (+) top, negative (-) bottom). Positive and negative response neurons increase and decrease their firing rate, respectively, upon sniffing onset. Total numbers of neurons are indicated at the upper left corner of each PSTH. Shading indicates ±SEM. **(E)** Venn diagram summary of target neuronal proportions for WT (left) and IRSp53-KO (right) pExc neurons (n = 233 [WT-pExc] and 258 [KO-pExc]). Numbers indicate neuronal proportion % (n neurons). E, empty, S, social, O, object. **(F)** Neuronal proportions for WT (black) and IRSp53-KO (red) positive and negative empty, social and object pExc neurons. Note that the majority of social target neurons respond positively to the social target. Note also that the IRSp53-KO social neuronal proportion is less than 50% (7/15) of the corresponding proportion for WT. **(G)** Normalized Δ firing rate between firing rate at the target sniffing zone versus that at the center zone for positive response target neurons (empty neuron, n = 6 [WT-pExc] and n = 9 [KO-pExc]; social neuron, n = 13 [WT-pExc] and n = 7 [KO-pExc]; object neuron, n = 9 [WT-pExc] and n = 8 [KO-pExc], **p<0.01, ns, not significant, two-way RM-ANOVA with Sidak’s multiple comparison’s test). See **Supplementary file 2** for statistics. Numerical data used to generate the figure are available in the **Figure 7—source data 1**. Figure 7—source data 1 Source files for target neuron data in Figure 7 The excel file contains the numberical data used to generate Figure 7A, B, F and G.

In order to determine whether the social, object, and empty neurons increase or decrease their firing rates upon target sniffing, we generated the average peristimulus time histograms (PSTHs) for WT and IRSp53-KO target neurons filtered by a z-score cut-off value of 1.0 (positive response neurons: z-score ≥ 1.0, negative response neurons: z-score ≤ -1.0). We found both positive and negative response target neurons (i.e., those increasing and decreasing their firing rates upon sniffing onset, respectively) in WT as well as IRSp53-KO mice (**Figure 7C,D, Figure 7 — figure supplement 1B**).

The numbers of target neurons were plotted in Venn diagrams (**Figure 7E**). The majority of social neurons were positive response neurons (**Figure 7F**), which is consistent with a previous report (Lee et al., 2016). In addition, the normalized Δ firing rate (i.e., target sniffing zone firing – center zone firing) of the positive target neurons was significantly lower in IRSp53-KO mice than WT mice (**Figure 7G**).

The decreases in the proportion of social-responsive pExc neurons and the response magnitude of target neurons may be associated with the social impairment seen in IRSp53-KO mice.

### Normal proportion of broadly-tuned target pExc neurons in the mPFC of IRSp53-KO mice

While the majority of target neurons were specific to a single target (single-tuned neurons), several target neurons were responsive to multiple targets (broadly-tuned neurons; **Figure 7E**). Examination of the PSTHs for examples of single-tuned and broadly-tuned neurons revealed that the target discriminability of WT and IRSp53-KO single-tuned neurons appeared to be higher than that of a subset of broadly-tuned neurons (**Figure 7—figure supplement 2A,B**). We found, however, that the overall proportions of single-tuned and broadly-tuned neurons were comparable between genotypes (**Figure 7—figure supplement 2C,D**).

## Discussion

The present study investigated abnormalities in social representation and neuronal firing patterns in the mPFC of IRSp53-KO mice. A key finding of the study is that social deficits in IRSp53-KO mice are associated with impaired social representation in the mPFC, which relates to a decreased firing rate variability and limited firing-rate range in IRSp53-KO putative excitatory neurons. The reduction in the firing-rate range is accompanied by a significant decrease in the response magnitudes to social and non-social targets and a limited discriminability of social and non-social cues.

Previous studies suggested that disruption of the excitation/inhibition balance (E/I imbalance) can cause abnormal social behaviors (Lee et al., 2017; Nelson and Valakh, 2015; Sohal and Rubenstein, 2019). This concept of an E/I imbalance can be applied to multiple mechanistic levels, ranging from synapses to neurons and neural circuits. Elevation of the E/I ratio, via optogenetic excitation of mPFC pyramidal neurons, was reported to impair social behavior in WT mice and decrease the synaptic current-mediated firing-rate ranges of neurons to suppress the dynamic range of information transfer (Yizhar et al., 2011). Here, we provide an in vivo example of this concept by demonstrating that IRSp53-KO mPFC excitatory neurons show increased spontaneous firing activity but a limited firing-rate range during social and non-social cue exploration.

The heightened firing rates of mPFC pExc neurons in IRSp53-KO mice during the rest period may be attributable to the increased intrinsic excitability of pyramidal mPFC neurons, as reported in IRSp53-KO mice with gene deletion restricted to excitatory neurons (Kim et al., 2020). An elevated firing rate at rest was observed for mPFC neurons in socially impaired *Cntnap2*-KO mice; this was correlated with a reduction in the signal-to-noise ratio and disruption of social sensory stimuli representation (Levy et al., 2019). Likewise, we herein report that IRSp53-KO mPFC excitatory neurons may also have ‘noisy’ properties that disrupt the reliable filtration and transduction of important signals, such as social cues.

Suppressed excitatory synaptic transmission accompanying reduced dendritic spine density and cognitive and social declines has been frequently observed in mouse models of neuropsychiatric disorders, including IRSp53-KO, Shank2-KO, Cntnap2-KO, and Syngap1-KO mice (Chung et al., 2015; Clement et al., 2012; Lazaro et al., 2019; Schmeisser et al., 2012). In our study, the limited firing-rate range of IRSp53 pExc neurons is expressed as reduced response magnitudes to social and non-social cues and a decreased ability to discriminate social and non-social cues. These insufficient firing-rate responses to external cues may be attributable to the cortical decreases in excitatory synapse density, dendritic spine density, and postsynaptic density (PSD) maturity seen in IRSp53-KO mice (Chung et al., 2015), which would substantially limit the amount of information delivered to and integrated at the mPFC under social contexts.

Dysfunctions of NMDA receptors have been associated with social deficits in mouse models of psychiatric disorders (Chung et al., 2019; Lee et al., 2021b; Lee et al., 2015; Mielnik et al., 2021; Shin et al., 2020; Won et al., 2012; XiangWei et al., 2018). In addition, IRSp53-KO mice show NMDAR hyperfunction and memantine treatment-dependent rescue of social deficits (Bobsin and Kreienkamp, 2015; 2016; Chung et al., 2015; Kim et al., 2009). Intriguingly, modulation of NMDAR activity has substantial influences on the firing rate, burst activity, and firing variability of PFC neurons in vivo (Homayoun and Moghaddam, 2006). In line with these findings, we herein report that IRSp53-KO mPFC pyramidal neurons show decreased burst firing and firing variability. It would be interesting to investigate whether memantine treatment of IRSp53-KO mice, which rescues social deficits, could also normalize the decreased burst and firing variability and the abnormal social representation in mPFC neurons.

The most salient feature observed herein for IRSp53-KO mPFC neurons is the reduced proportion of social neurons, but not those for other targets, that is consistently observed across all z-score cut-off ranges. Disruptions in the cortical social representation of social contexts have been reported in several studies of ASD models (Cao et al., 2018a; Lee et al., 2021a; Lee et al., 2021b; Levy et al., 2019). The present study further highlights the importance of a reduced social/non-social encoding neuron ratio in social deficits. It should be pointed out, however, that it remains unclear whether the limited social cortical representation that is strongly associated with social deficits represents the cause of limited social brain functions, versus being an outcome of reduced social interaction or even limited sensory input. The abovementioned pharmacological rescue experiments attempting to correct both cortical social representation and social behaviors might help clarify this causal relationship.

In summary, our current results indicate that IRSp53-KO mice display elevated spontaneous firing and reduced firing variability and range in mPFC neurons, which may suppress cortical social representation and induce behavioral social deficits.

## Materials and Methods

### Animals

Adult (3–5 months old) C57B/6J male WT (n = 6) and IRSp53-KO (n = 8) mice were used for single-unit recording. Mice were fed *ad libitum* and maintained under 12-h light/dark cycle (light period 1 am–1 pm). All experiments were conducted during the dark phase (1 pm–1 am) of the light/dark cycle. The mouse facility and experimental setting were always maintained at 21°C and 50–60% humidity. Mice were maintained according to the Animal Research Requirements of Korea Advanced Institute of Science and Technology (KAIST). All experiments were conducted with approval from the Committee on Animal Research at KAIST (approval number KA2020-94).

### Linear-chamber social-interaction test

Before linear-chamber social-interaction test (Lee et al., 2017), the subject mouse was placed in a white 7.5 cm radius x 15 cm opaque acryl container for neural recording at rest. The mouse was then allowed to explore the 45 cm x 10 cm x 21 cm linear chamber with the empty-empty chambers (E-E session), the social-object chambers (first S-O session), and then the same object-social chambers with the side exchanged (second S-O session), for 10 minutes each. The social and object targets used for each experiment were always novel. Novel male 129/Sv mice of similar age were used as the social target. The initial placement of social and object targets into left or right chambers in the first S-O session were randomly chosen before each experiment. All recordings were conducted at 30 lux. A total of 12 recording experiments were conducted for each mouse, with 3 days of isolation interval.

#### Mice movement tracking

Mice movements were monitored by a digital camera mounted on the ceiling, directly above the linear chamber assay. The position of the mouse’s nose, right and left ears, and tail-base, and four corner points of the linear chamber were trained using a pose estimation software DeepLabCut (version 2.0; Mathis et al., 2018).

#### Behavioral analysis

Sniffing time was defined as the time when the nose point is within 3 cm from the face of each target chamber. In-zone time was defined as the time when the body center (midpoint between nose point and tail base) is within 9 cm from the face of each target chamber.

### Single-unit recording

Eight tetrodes were implanted in the mPFC (four tetrodes per hemisphere; 1.7–2.1 mm anterior and 0.1–0.5 mm lateral from bregma, and 1.5–2.3 mm ventral from brain surface). 36-channel electrode interface board (EIB-36; Neuralynx, Bozeman, MT, USA) and hyperdrive (modified version of Flex drive from Open Ephys) were used. Mice were subjected to 3 days of handling (10 min each day) after 1 week of recovery from surgery. At the first exposure to the linear chamber test, mice were habituated to the environment and tether but without recording. After habituation, mice were subjected to 12 linear-chamber experiments with recording. 10,000x amplified single-unit recording signals with 32kHz sampling frequency were filtered using a bandpass filter of 600–6000 Hz. Signals were recorded via Digitalynx (hardware; Neuralynx, Bozeman, MT, USA) and Cheetah data-acquisition system (software version 5.0; Neuralynx, Bozeman, MT, USA) and stored in a personal computer. In order to record different units at each recording experiment, the positions of tetrodes were lowered by 62.5 µm after the recording.

#### Histology

After the 12th recording, mice were deeply anesthetized and the locations of the tetrodes were marked by electrolytic lesion (100 μA unipolar current for 7 sec for each electrode) and brains were extracted and perfused in 4% Paraformaldehyde (PFA) solution for at least 72 hours. The fixed brains were sliced coronally (50 µm) using a vibratome (VT1000; Leica, Buffalo Grove, IL, USA), stained with DAPI, and the positions of lesions were assessed by post hoc histological evaluation using a confocal microscope (LSM780; Carl Zeiss, Oberkochen, Germany).

### Spike analysis

The single-unit spike clusters were isolated manually by spike waveform features, such as, energy, peak, valley, and principal components, using MClust (version 4.4, available online at http://redishlab.neuroscience.umn.edu/mclust/MClust.html; credits to A. David Redish). Only units with isolation distance of ≥ 25 and L-ratio of ≤ 0.1 were used for analysis.

Only valid sniffing trials and valid in-zone (and center zone) trials were used for spike analysis. Valid sniffing trials were defined as those with a duration of ≥ 1 sec and inter-trial interval of ≥ 2 sec. For sufficient acquisition of center zone trials, valid in-zone (and center zone) trials were defined as those with a duration of ≥ 0.5 sec and inter-trial interval of ≥ 0.5 sec. Neurons with missing valid sniffing and in-zone trials for any of the six targets (left and right for the E-E session, social and object for the first and second S-O sessions) were excluded from spike analysis. The number of neurons recorded after the valid trial exclusion was WT n = 391 total neurons, 366 pExc neurons, 17 pInh neurons from 6 mice and IRSp53-KO n = 394 total neurons, 359 pExc neurons, 24 pInh neurons from 8 mice (see Supplementary file 1 for details). The pExc and pInh neurons were classified based on half-valley width (pExc: HVW > 200 ms, pInh: HVW < 200 ms) and peak-to-valley ratio (pExc: PVR > 1.4, pInh: PVR < 1.4).

Except for the comparison of mean firing rate at rest (total number of spikes within 5-min resting duration), only pExc neurons with the average firing rate of ≥0.5 Hz during the 30-min linear chamber assay were used for further analysis. After ≥0.5 Hz filtration, the total number of pExc neurons were WT n = 233 neurons from 6 mice and IRSp53-KO n = 258 neurons from 8 mice (see Supplementary file 1 for details).

#### Instantaneous firing rate and firing rate variability analysis

For instantaneous firing rate analysis, the 30-min linear chamber period were divided into 1800 time-bins (3-sec time-bin of 1-sec steps). The sigma (Hz) value for each neuron was defined as the 1 standard deviation (1SD; includes 68% of data) value of the 1800 instantaneous firing rates. The normalized instantaneous firing rate of each neuron was calculated by dividing the instantaneous firing rates by the maximum instantaneous firing rate.

#### ISI and Burst analysis

Interspike interval (ISI) is the time between two consecutive spikes (in ms). For the average ISI histogram, ISIs ≤ 200 ms were extracted for each neuron, and the ISI histogram values of individual neurons were averaged. The coefficient of variation (CV) of ISI for each neuron was calculated as σ_ISI_/μ_ISI_, in which σ_ISI_ is the 1SD of ISI and μ_ISI_ is the mean of ISI.

Burst proportion (%) was defined as the number of burst spikes out of the total number of spikes in a neuron, in which the burst spikes are defined as all consecutive spikes with an ISI ≤ 10 ms. To demonstrate the cut-off effects of ISI burst definition, burst analysis was performed using a range of burst ISI threshold values (5 – 30 ms).

To compare the composition of the burst events between genotypes, all burst events were classified into doublet, triplet and ≥4 spike groups according to the number of spikes in individual burst events. The proportions of burst events by spike count were then compared between genotypes via Chi-square analysis.

#### Maximum Δ firing rate analysis

The maximum Δ firing rate of a neuron is the maximum value between the absolute firing rate differences between the firing rates at the two in-zones and the firing rate at the center zone (|FR_I1_ – FR_C_| and |FR_I2_ – FR_C_|) where FR_c_ is the firing rate at the center zone and FR_I1_ and FR_I2_ are the firing rates at two in-zones). The two in-zones are left and right for the E-E session, and social and object for the first and second S-O sessions.

The normalized maximum Δ firing rate is the maximum value between absolute normalized firing rate differences between the firing rates at two in-zones and the firing rate at the center zone (|(FR_I1_ – FR_C_)/(FR_I1_ + FR_C_)| and |(FR_I2_ – FR_C_)/(FR_I2_ + FR_C_)|).

#### Discrimination index analysis

The neuron’s discriminability between targets was assessed by calculating the discrimination index, which is the normalized firing rate differences between the firing rates at the two in-zones: (FR_L_ – FR_R_)/(FR_L_ + FR_R_) for the E-E session and (FR_S_ – FR_O_)/(FR_S_ + FR_O_) for the S-O sessions where FR_L_, FR_R_, FR_S_, and FR_O_ are the mean firing rate at left, right, social, and object in-zones, respectively.

#### Target (empty, social, object) neuron analysis

For each neuron, the average firing rate during the valid sniffing time of empty (E), social (S), and object (O) targets were calculated. Its center zone time in the E-E, first S-O, and second S-O sessions were extracted, divided into 0.5-sec time-bins, and its instantaneous firing rates were calculated. The z-scores for each neuron were defined as (FR_T_ - µ_C_)/σ_C_, in which FR_T_ is the mean firing rate during target (E, S, or O) sniffing, while µ_C_ and σ_C_ are the mean and 1SD of the instantaneous firing rates at the center zone, respectively.

A range of z-score thresholds (0 – 2.58) was used to determine the neurons that are responsive to E, S, O targets. The proportions of WT and IRSp53-KO pExc target neurons (the number of target neurons out of the number of total neurons (in %)) were considered statistically different if the nonlinear fitted lines (third-order polynomial fit; across all calculated neuronal proportions for 0 – 2.58 z-score threshold range) were significantly different (via comparison of fits in Prism 9.0; GraphPad, San Diego, CA, USA). Z-score threshold value of 1.0 (positive response neuron: z-score ≥ 1.0, negative response neuron: z-score ≤ 1.0) was used for generating peristimulus time histograms (PSTHs) and comparing the normalized Δ firing rates of target neurons.

For PSTH of an individual neuron, firing rates were calculated in 250-ms bins (from -1.5 sec to 3 sec after the onset of target sniffing) and averaged across the sniffing trials. For the averaged PSTH of target neurons, the averaged firing rates of individual neurons were normalized by their maximum firing rate, and then averaged across all target neurons.

The normalized Δ firing rate of positive response target neurons is the normalized difference between the firing rates at the target zone (empty, social, or object depending on which target the target neuron is responsive to) and the center zone: (FR_T_-FR_C_)/(FR_T_+FR_C_) where FR_C_ is the firing rate at the center zone and FR_T_ is the firing rate at the target zones. Target zones fall in the area within 9 cm from the front face of the target chamber (same as in-zones).

### Statistical analysis

Statistical significance was determined via repeated measures of two-way ANOVA with Sidak’s multiple comparisons test (or Bonferroni’s multiple comparisons test), Friedman test with Dunn’s multiple comparisons test, Mann-Whitney test, simple linear regression with slope comparisons test, nonlinear fit with comparisons of fits, and Chi-square test (all via Prism 9.0; GraphPad, San Diego, CA, USA). Kolmogorov-Smirnov normality test was used to determine whether to use a parametric or nonparametric test. Graphs were generated by MATLAB 2020a (MathWorks, Natick, MA, USA) and Prism 9.0. All box and whisker plots show median, interquartile range, and 2.5 and 97.5 percentile. See **Supplementary file 2** for details on statistics.

## Supplementary file 1

Table 1. Recorded neuron number for each mouse

## Supplementary file 2

Details on the statistical tests and p-value.

## Acknowledgments

We would like to thank Changho Jo and Seohui Bae for their help with DeepLabCut. This study was supported by the National Research Foundation of Korea (NRF) grants (No. NRF-2019R1A2C4069863 to S.P.), a faculty research grant of Yonsei University College of Medicine (6-2020-0089 to E.L.), Korea Research Foundation (NRF-2019R1A2C1084812 to E.L.), IBS-R002-D2 (to M.W.J.), and IBS-ROO2-D1 (to E.K.). Figure 1A, 1C, 2C, and 5A were created with BioRender.com.

## Competing interests

The authors declare no competing financial interests.

---

**Figure 1–figure supplement 1.**
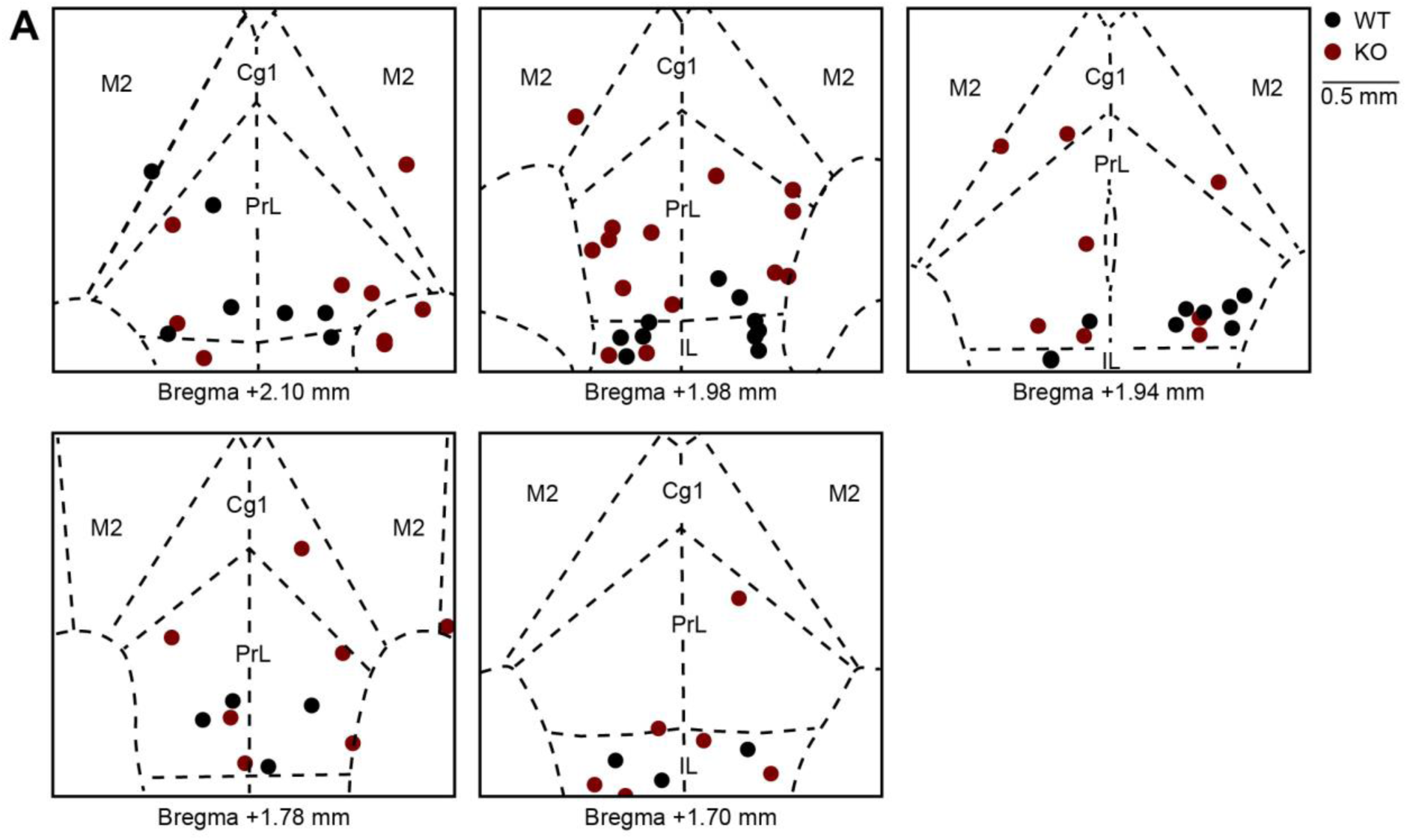
Locations of implanted tetrodes in the mPFC of WT and IRSp53-KO mice. **(A)** Final locations of tetrodes implanted into the mPFC of WT (black) and IRSp53-KO (red) mice. Coronal sections of the mPFC shown in this figure represent +1.70–+2.10 mm away from the bregma. PrL, prelimbic cortex; IL, infralimbic cortex; Cg1, cingulate cortex, area 1; M2; secondary motor cortex.

**Figure 1–figure supplement 2.**
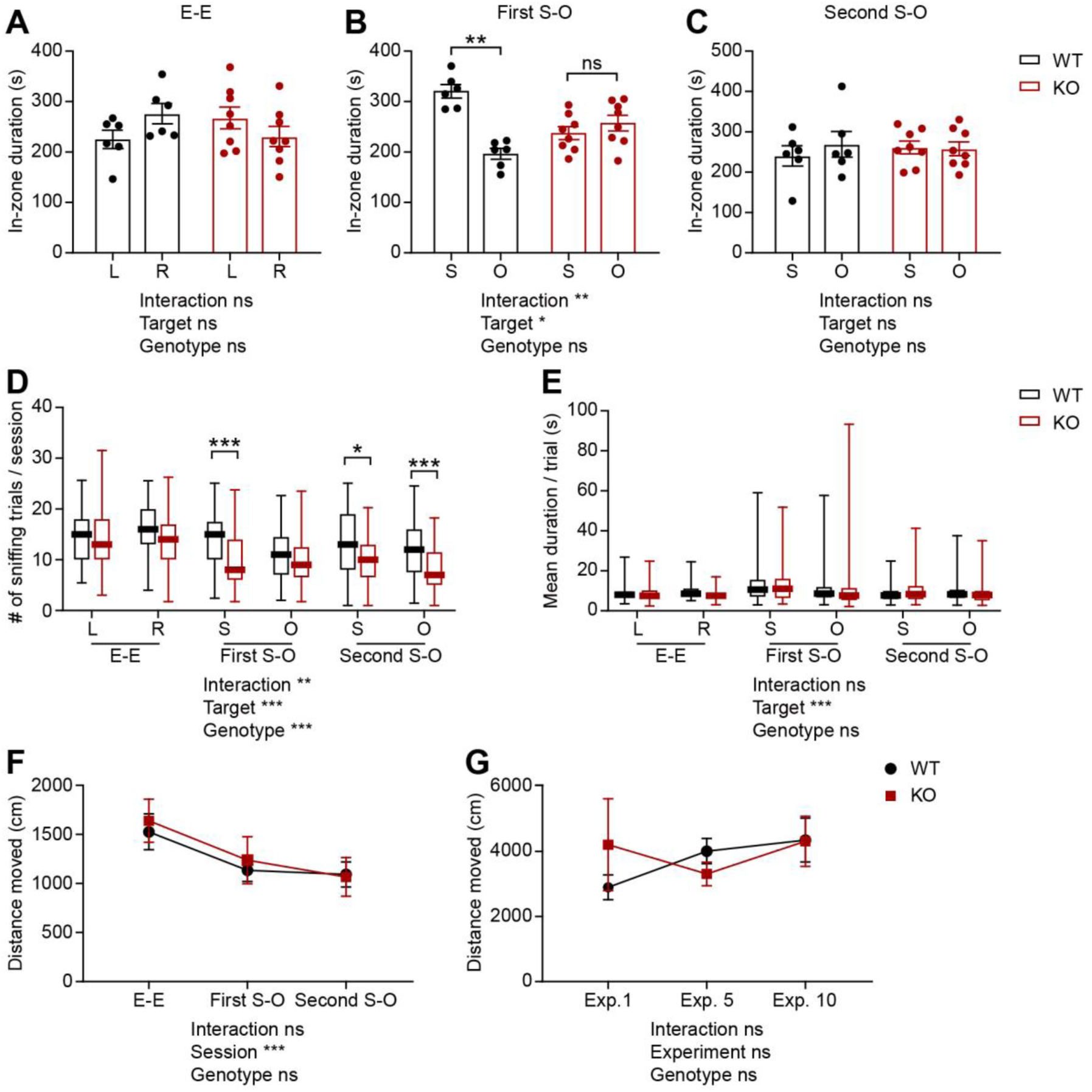
Social impairments, as judged by in-zone durations, and unaltered mean duration of each visit or locomotor activity in IRSp53-KO mice in the linear social-interaction chamber. **(A–C)** Mean in-zone durations for left (L) vs. right (R) empty targets during the E-E session and social (S) vs. object (O) targets during the first and second S-O sessions. (n = 6 mice [WT], 8 mice [IRSp53-KO], *p < 0.05, **p < 0.01, ns, not significant, two-way RM-ANOVA with Sidak’s multiple comparison test). **(D and E)** The average number of sniffing visits (**D**) and mean duration of time spent sniffing per valid sniffing trial (**E**) for each target during the E-E, first S-O, and second S-O sessions. (n = 57 experiments from 6 mice [WT], 69, 8 [IRSp53-KO], *p < 0.05, **p < 0.01, ***p < 0.001, ns, not significant, two-way RM-ANOVA with Sidak’s multiple comparison test). **(F and G)** The average (±SEM across 6 WT mice and 8 IRSp53-KO mice) distance moved in the linear social-interaction test across three consecutive sessions (E-E, first S-O, and second S-O) in an experiment (**F**) and across different recording experiments (1st, 5th, and 10th experiments used as examples; **G**). (n = 6 mice [WT], 8 mice [IRSp53-KO], ***p < 0.001, ns, not significant, two-way RM-ANOVA with Sidak’s multiple comparison test). See **Supplementary file 2** for statistics. Numerical data used to generate the figure are available in the **Figure 1—figure supplement 2—source data 1**. Figure 1—figure supplement 2—source data 1 **Source files for mouse behavior data in Figure 1—figure supplement 2** The excel file contains the numberical data used to generate Figure 1—figure supplement 2A–G.

**Figure 3–figure supplement 1.**
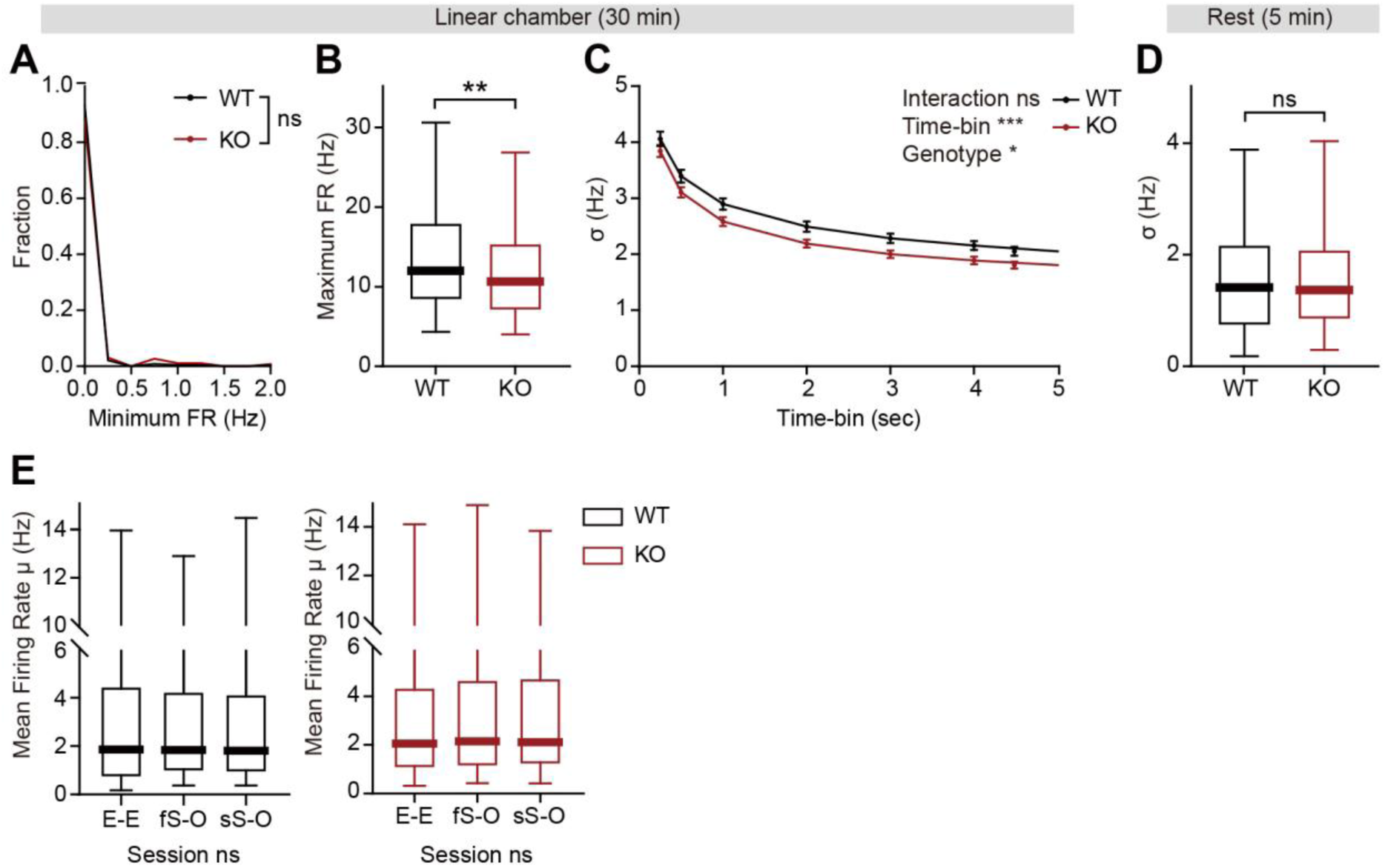
Decreased firing-rate variability in IRSp53-KO pExc mPFC neurons selectively during linear chamber exploration. **(A and B)** Minimum (**A**) and maximum (**B**) instantaneous firing rates of WT and IRSp53-KO pExc neurons during the 30-min linear chamber test. (n = 233 [WT-pExc] and 258 [KO-pExc], **p < 0.01, Mann-Whitney test). **(C)** Average (±SEM) sigma values of instantaneous firing rates during the 30-min linear chamber test calculated using different time-bin sizes (0.25–5 sec). Window sizes were set to be the same as the time-bin size. Note that the overall sigma values of IRSp53-KO neurons are significantly smaller than those of the WT neurons for all analyzed time-bin sizes. (n = 233 [WT-pExc] and 258 [KO-pExc], *p < 0.05, ***p < 0.001, ns, not significant, two-way RM-ANOVA). **(D)** Sigma values for the instantaneous firing rates (3-sec window advanced in 1-sec steps) during the 5-min rest period in WT and IRSp53-KO pExc neurons. (n = 233 [WT-pExc] and 258 [KO-pExc], ns, not significant, Mann-Whitney test). **(E)** Mean instantaneous firing rates of WT (left) and IRSp53-KO (right) pExc neurons during the E-E, first S-O, and second S-O sessions of the linear chamber test. (n = 233 [WT-pExc] and 258 [KO-pExc], ns, not significant, Friedman test). See **Supplementary file 2** for statistics. Numerical data used to generate the figure are available in the **Figure 3—figure supplement 1—source data 1**. **Figure 3—figure supplement 1—source data 1** **Source files for instantaneous firing rate data in Figure 3—figure supplement 1** The excel file contains the numberical data used to generate Figure 3—figure supplement 1A–E.

**Figure 4–figure supplement 1.**
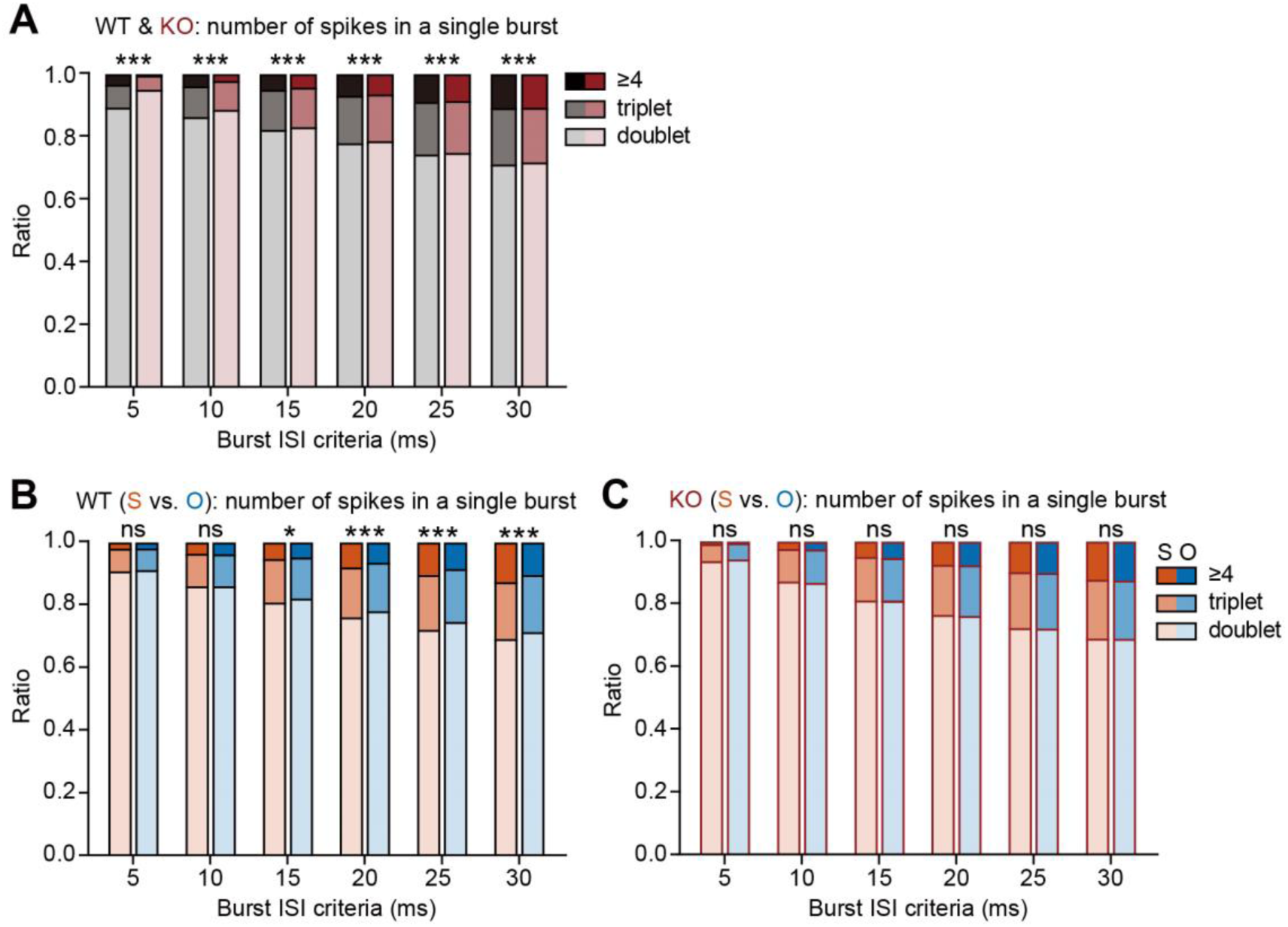
Lower discriminability between social and object targets by compositions of burst events in IRSp53-KO pExc mPFC neurons. **(A)** Numbers of spikes in each burst (doublet, triplet, and ≥4) for different burst ISI thresholds. All burst events from entire recordings of 233 WT pExc neurons and 258 IRSp53-KO pExc neurons were used for analysis. Note that as the burst ISI threshold increases, the proportion of bursts with three or more consecutive spikes (triplet and ≥4 groups) increases. Note also that most of the bursts (∼70 – 90%) are doublets. (***p<0.001, Chi-square test). **(B and C)** Numbers of spikes in each burst during social (S) versus object (O) sniffing for different burst ISI thresholds. All burst events recorded from 233 WT pExc neurons (**B**) and 258 IRSp53-KO pExc neurons (**C**) during social and object sniffing were used for analysis. (*p < 0.05, ***p < 0.001, ns, not significant, Chi-square test). See **Supplementary file 2** for statistics. Numerical data used to generate the figure are available in the **Figure 4—figure supplement 1—source data 1**. Figure 4—figure supplement 1—source data 1 **Source files for burst event composition data in Figure 4—figure supplement 1** The excel file contains the numberical data used to generate Figure 4—figure supplement 1A–C.

**Figure 6–figure supplement 1.**
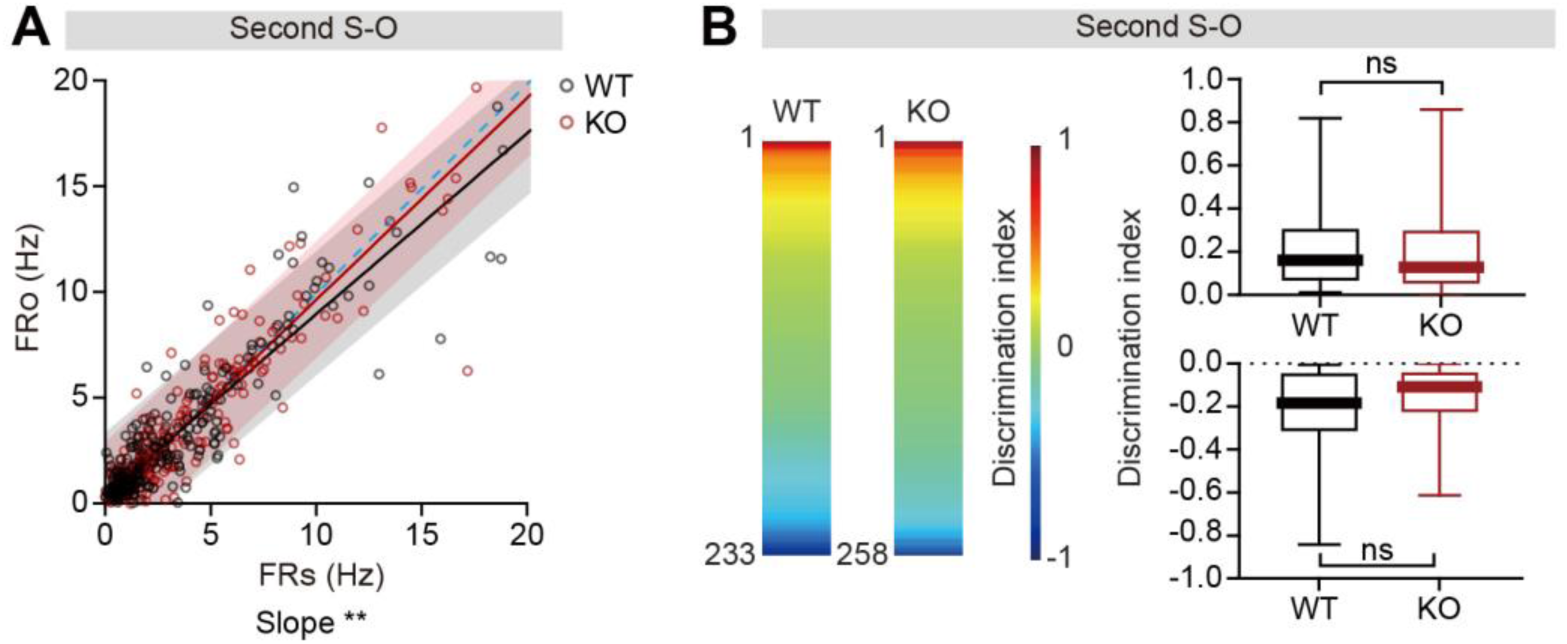
Normal firing-rate discriminability between social and object targets in IRSp53-KO pExc mPFC neurons in the second S-O session. **(A)** Scatterplot of social in-zone firing rate (FR_S_) against object in-zone firing rate (FR_O_) during the second S-O session for WT and IRSp53-KO pExc neurons. Solid lines indicate simple linear regressions for WT (black) and KO (red) neurons. Shaded areas indicate the 95% confidence intervals for WT (black) and KO (red) firing rates. Blue dashed lines are 45-degree lines. (n = 233 [WT-pExc] and 258 [KO-pExc], **p < 0.01, simple linear regression with slope comparison test). **(B)** Heatmaps (left) showing the discrimination index representing social and object target discriminability during the second S-O session for individual WT and IRSp53-KO pExc neurons sorted from 1 to –1 (n = 233 [WT-pExc] and 258 [KO-pExc]). Positive (top, n = 114 [WT-pExc] and n = 134 [KO-pExc]) and negative (bottom, n = 119 [WT-pExc] and n = 124 [KO-pExc]) discrimination indexes (right) represent pExc neurons with social > object and social < object discriminability, respectively, during the second S-O session. (ns, not significant, Mann-Whitney test). See **Supplementary file 2** for statistics. Numerical data used to generate the figure are available in the **Figure 6—figure supplement 1—source data 1**. Figure 6—figure supplement 1—source data 1 **Source files for firing-rate discriminability data in Figure 6—figure supplement 1** The excel file contains the numberical data used to generate Figure 6—figure supplement 1A–B.

**Figure 7–figure supplement 1.**
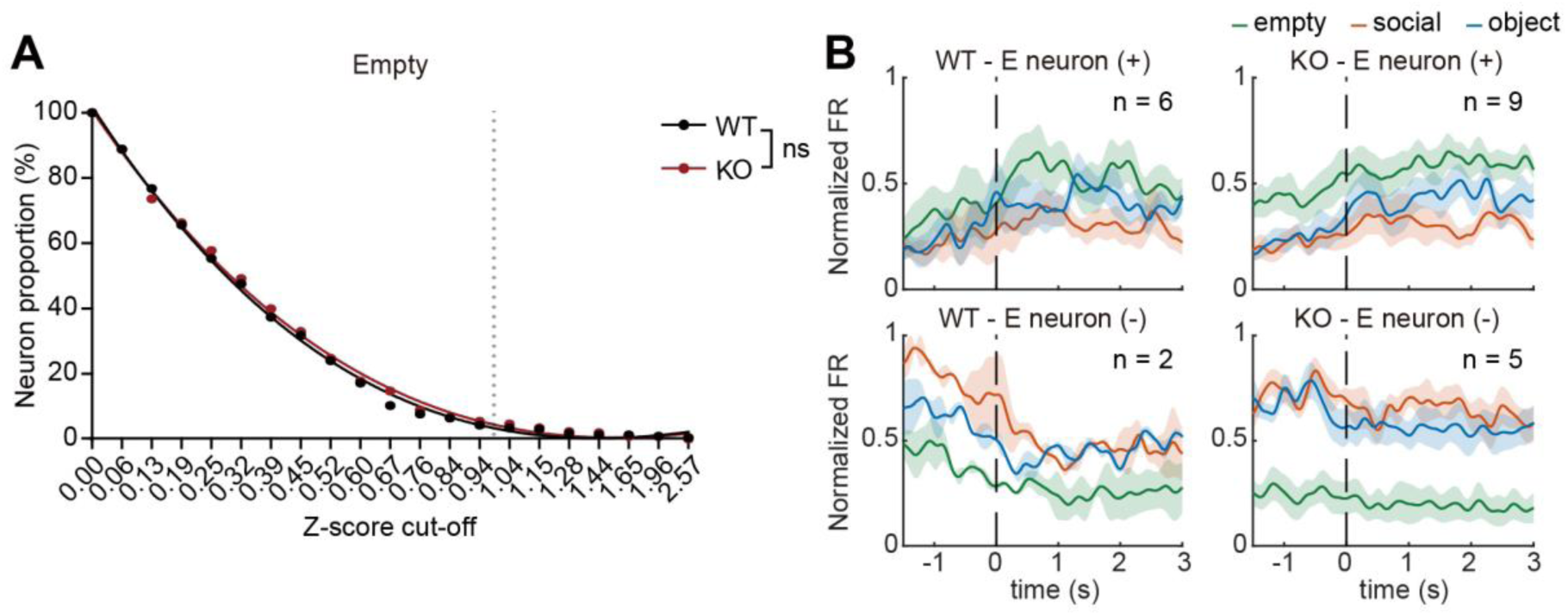
Normal proportion of empty pExc neurons in the mPFC of IRSp53-KO mice. **(A)** Empty neuronal proportions for 233 WT pExc neurons and 258 IRSp53-KO pExc neurons using a z-score cut-off range of 0 to 2.58. For each neuron, the mean z-scores of firing rates obtained during empty sniffing were normalized by the firing rates obtained at the center zone. See Methods for details on z-score calculation. Solid lines indicate the nonlinear fitted lines for the WT (black) and IRSp53-KO (red) groups. Dotted lines indicate z-score cut-off value of 1.0. (ns, not significant, comparison of nonlinear fits (see Methods)). **(B)** Average PSTHs of firing rate responses to empty (green), social (orange), and object (blue) targets (aligned to the onset of sniffing) for all empty pExc neurons filtered by a z-score cut-off value of 1.0. Empty neurons are divided by genotype (WT left, IRSp53-KO right) and response direction (positive (+) top, negative (-) bottom). Positive and negative response neurons increase and decrease their firing rate, respectively, upon sniffing onset. Total numbers of neurons are indicated at the upper left corner of each PSTH. Shading indicates ±SEM. See **Supplementary file 2** for statistics. Numerical data used to generate the figure are available in the **Figure 7—figure supplement 1—source data 1**. Figure 7—figure supplement 1—source data 1 **Source files for empty neuron data in Figure 7—figure supplement 1** The excel file contains the numberical data used to generate Figure 7—figure supplement 1A.

**Figure 7–figure supplement 2.**
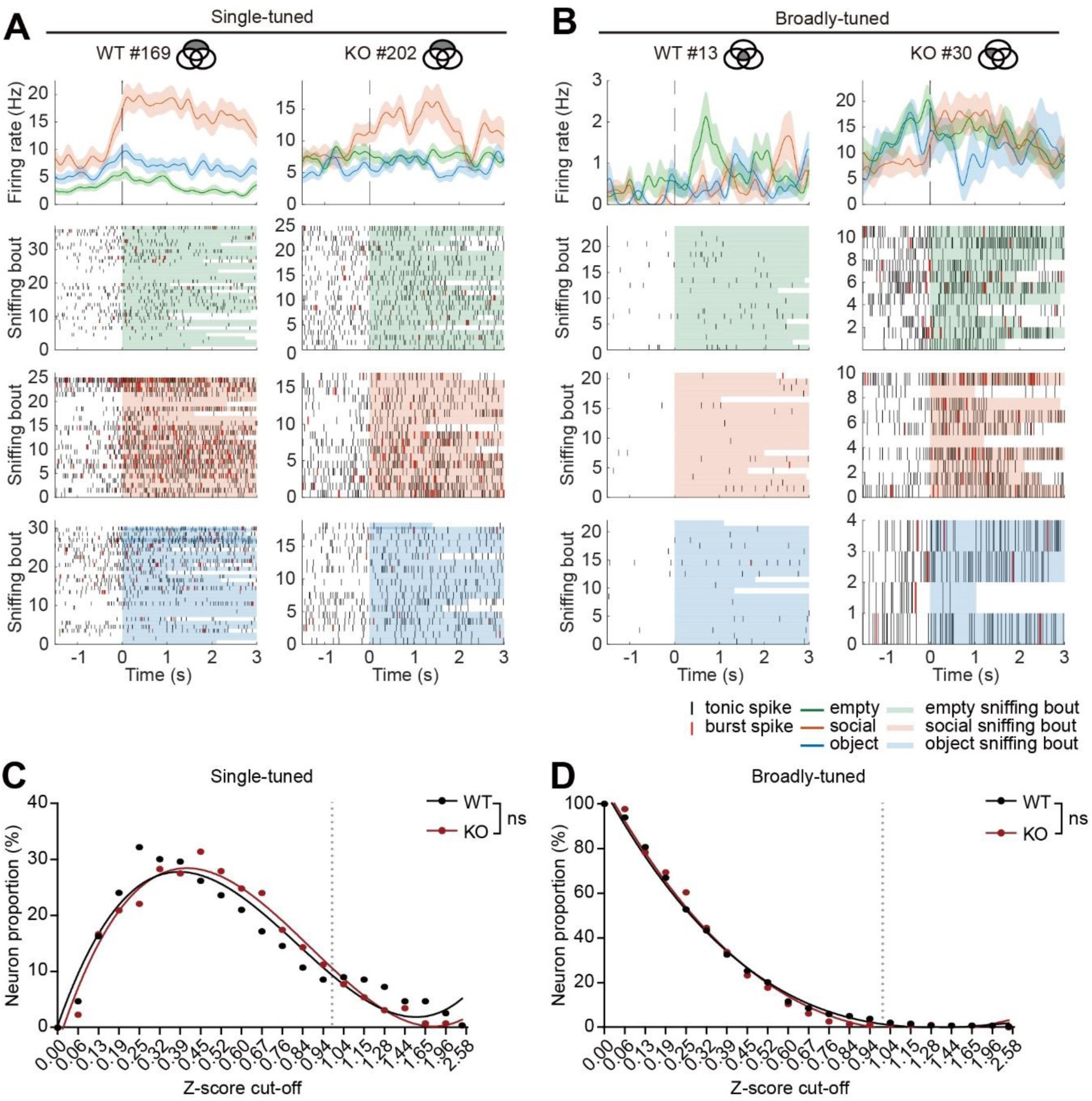
Comparable proportions of single-tuned and broadly-tuned pExc neurons in mPFC of IRSp53-KO mice. **(A and B)** Average PSTHs (top) and spike raster plots (bottom) of target sniffing responses (aligned to the onset of sniffing) for a single-tuned (**A**) and a broadly-tuned (**B**) WT social (left) and IRSp53-KO social (right) example pExc neurons. Single-tuned and broadly-tuned neurons are at the non-overlapping and overlapping regions of the Venn diagram (Figure 7E), respectively. Shading in PSTH indicates ±SEM across sniffing trials. **(C and D)** Single-tuned (**C**) and broadly-tuned (**D**) neuronal proportions obtained from 233 WT pExc neurons and 258 IRSp53-KO pExc neurons using a z-score cut-off range of 0 to 2.58. Solid lines indicate nonlinear fitted lines for the WT (black) and IRSp53-KO (red) groups. Dotted lines indicate z-score cut-off value of 1.0. (ns, not significant, comparison of nonlinear fits (see Methods)). See **Supplementary file 2** for statistics. Numerical data used to generate the figure are available in the **Figure 7—figure supplement 2—source data 1**. Figure 7—figure supplement 2—source data 1 **Source files for target neuron data in Figure 7—figure supplement 2** The excel file contains the numberical data used to generate Figure 7—figure supplement 2C and D.

## Notes

**Competing interests:** Eunjoon Kim: Reviewing editor, *eLife*

### Competing Interest Statement

The authors have declared no competing interest.

